# The optical origin of the human skin color ‘banana’ in CIELAB space

**DOI:** 10.64898/2026.06.16.732713

**Authors:** Maysoon Harunani, Ye Jin Han, Mutian Shen, Blake Sparkman, David Chen, Zohar Nussinov, Leonid Shmuylovich

**Affiliations:** Department of Biomedical Engineering, Washington University in Saint Louis, MO 63110; Department of Physics, Washington University in Saint Louis, MO 63110; Department of Medicine, Division of Dermatology, Washington University in Saint Louis, MO, 63110

**Keywords:** Tissue optics, colorimetry, objective skin color measurement, adding doubling, Monte Carlo, Individual Typology Angle

## Abstract

Human skin colors occupy a characteristic banana-shaped region in CIE L*a*b* space, but why skin color coordinates are restricted to this region and how they relate to melanin and blood remain incompletely understood. We developed a physics-based framework linking skin chromophore content to colorimeter-derived skin color coordinates using two complementary three-layer light transport models. Across physiologic ranges of epidermal melanosome volume fraction and dermal blood volume fraction, simulated reflectance spectra were converted to CIE L*a*b* coordinates and compared with human skin color measurements from the International Skin Spectra Archive. Physiologic variation in melanin and blood reproduced the observed banana-shaped locus and revealed distinct chromophore-specific trajectories. Iso-melanin trajectories became progressively more linear as melanin increased, whereas iso-blood trajectories retained the curvature of the skin color locus. As melanin increased, perceptible color differences from blood volume changes were reduced, providing a mechanistic explanation for reduced erythema visibility in highly pigmented skin. These relationships were stable across plausible variations in layer thickness and tissue oxygenation and agreed with external validation data. The framework also identified when the Individual Typology Angle is confounded by blood or distorted by dermal melanin. Together, these findings establish a mechanistic optical basis for interpreting colorimeter-derived skin color coordinates.

## INTRODUCTION

Skin comprises multiple layers, each containing distinct absorbing and scattering constituents that shape its optical appearance. At visible wavelengths, the dominant chromophores are hemoglobin and melanin (Zonios et al., 2001), and disease-associated changes in these components lead to recognizable alterations in skin color. For example, increases in dermal blood volume produce erythema, a key clinical sign of inflammation. However, epidermal melanin strongly absorbs visible light at wavelengths used to visualize hemoglobin and can therefore make erythema less perceptible in highly pigmented skin. These same chromophores also influence the performance of optical medical devices. For example, pulse oximeters have been reported to be less accurate in darkly pigmented individuals, and melanin-dependent optical interactions have been implicated in this disparity (Feiner et al., 2007). Therefore, in both clinical care and device development, there is a strong interest in noninvasively quantifying skin melanin and hemoglobin to improve diagnosis, guide management, and ensure equitable device performance.

While diffuse reflectance spectroscopy has been explored as a method for assessing skin chromophores (Pryor et al., 2025; Ly et al., 2020), objective skin color assessment in clinical settings is primarily performed with colorimeters, which measure skin reflectance and map it into CIELAB color space. Each measurement yields a coordinate in this space, commonly visualized in the L*–b* plane, where L* represents lightness and b* represents the blue-yellow chromatic axis (Figure 1A). CIELAB is an opponency color space with three components: lightness (L*) and two chromatic axes, a* (green to red) and b* (blue to yellow) (Chardon et al., 1991). Importantly, unlike device-dependent RGB values, CIELAB was designed so that the Euclidean distance between coordinates, ΔE, approximates perceptual color difference (Ly et al., 2020). In landmark studies spanning several decades (Chardon et al., 1991; Del Bino et al., 2006; Del Bino et al., 2013), investigators measured CIELAB skin color coordinates in thousands of individuals and found that, when plotted in the L*-b* plane, human skin colors cluster within a characteristic ‘banana-shaped’ region, initially termed the “skin color volume”(Figure 1A) (Chardon et al., 1991).

**Figure 1.**
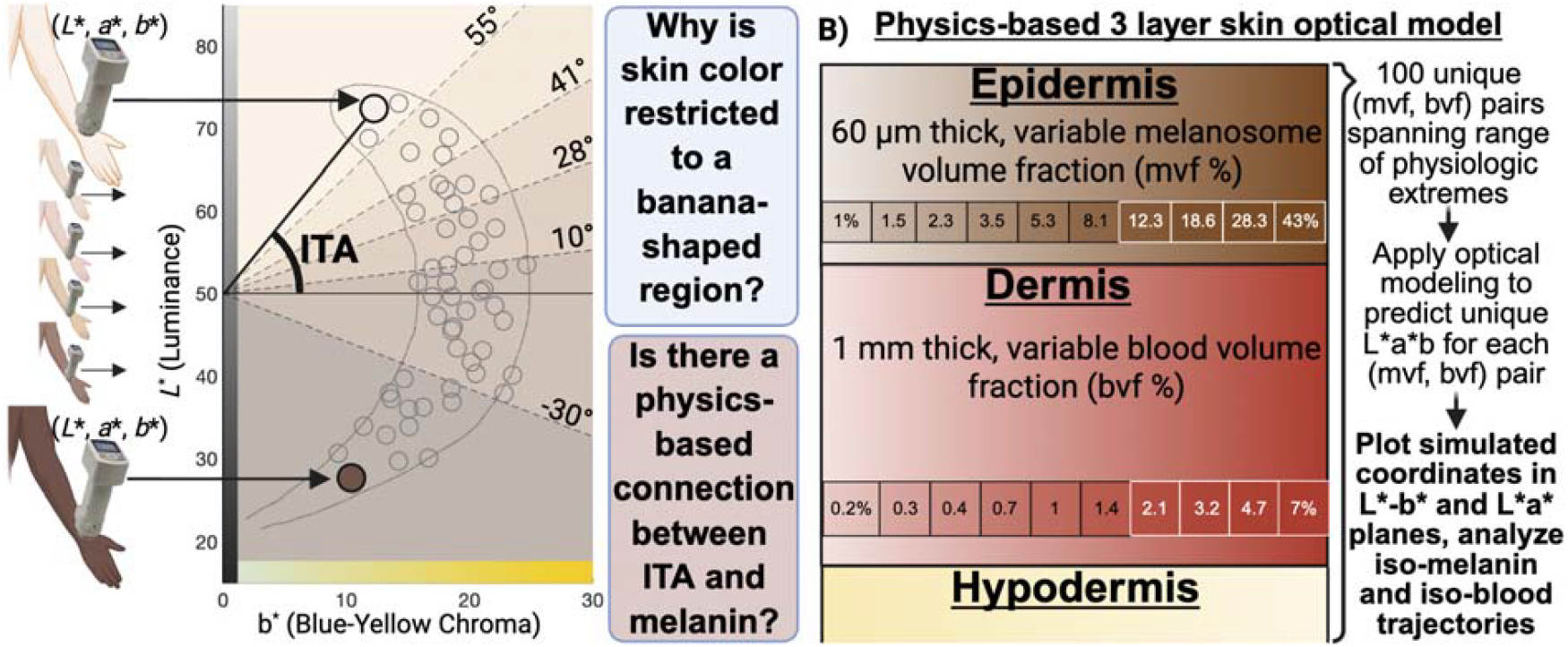
(A) Colorimetry maps measured skin reflectance to CIE L*a*b* coordinates, which can be visualized as points in the L*–b* plane. Population-level measurements occupy a characteristic banana-shaped skin color region, with lightly pigmented skin generally located toward higher L* values and darkly pigmented skin toward lower L* values. The Individual Typology Angle (ITA) is defined as the angle formed between a measured L*–b* coordinate and the reference point (b* = 0, L* = 50), with commonly used thresholds dividing the plane into six pigmentation categories. This schematic highlights two central questions addressed in this study: why skin color coordinates are restricted to a banana-shaped region, and whether a physics-based relationship connects ITA to underlying melanin and blood content. (B) Physics-based three-layer skin optical model consisting of a melanosome-containing epidermis, blood-containing dermis, and hypodermis. Melanosome volume fraction and blood volume fraction were varied logarithmically across physiologic ranges to generate 100 unique chromophore combinations. For each combination, optical modeling was used to predict reflectance spectra, convert spectra to L*a*b* coordinates, and define iso-melanin and iso-blood trajectories in color space.

Within this region, lightly pigmented skin occupies the upper portion of the curve, while darkly pigmented skin clusters toward the lower portion. Despite its reproducibility across populations, the shape and boundaries of this region remain empirical observations rather than predictions derived from skin physiology and optics. In particular, it remains unclear why human skin L*-b* coordinates are confined to this region, and which physiological parameters determine its limits.

Beyond defining the permissible range of skin color L*a*b* coordinates, investigators have also explored whether colorimeter outputs can reveal underlying chromophore content. Chardon et al. proposed estimating pigmentation with the Individual Typology Angle (ITA), defined as the angle subtended between the point (b*=0, L*=50) and a subject’s (b*, L*) coordinates (Figure 1A):

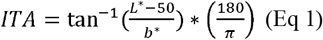

ITA thresholds are commonly used to divide subjects into categorical pigmentation groups, and many studies have reported an approximately linear relationship between ITA and melanin content (Chardon et al., 1991; Del Bino et al., 2006; Del Bino and Bernerd, 2013). However, a causal relationship between ITA and melanin concentration has not been established, and several groups have raised concerns that ITA is confounded by blood content (Stamatas and Kollias, 2004; Park et al., 1999; Takiwaki et al., 2002; Hou and Li, 2024). For example, Stamatas and Kollias showed that venous congestion induced by blood pressure cuff occlusion produces large changes in ITA despite unchanged melanin (Stamatas and Kollias, 2004). A second potential limitation is that ITA was developed from ordinary constitutive skin-color coordinates, which typically fall in the positive b* region, and may behave differently in dermatologic lesions where pigment depth, inflammation, hemorrhage, or vascular change alters the relationship between color and melanin. Together, these observations illustrate two key points: (i) L*a*b* coordinates shift with physiological changes in blood volume, and (ii) ITA may not be a specific surrogate for melanin. More broadly, empirical perturbation studies (hyperpigmentation through UV exposure, depigmentation from vitiligo, steroid-induced blanching, venous congestion) have mapped L*a*b* shifts with melanin and blood variation, but these relationships remain correlational rather than mechanistic (Chardon et al., 1991; Del Bino et al., 2006; Del Bino and Bernerd, 2013; Stamatas and Kollias, 2004; Park et al., 1999; Takiwaki et al, 2002; Hou and Li, 2024; Alaluf, et al., 2002).

These observations highlight two fundamental gaps, illustrated schematically in Figure 1A. First, although population-level colorimetry measurements occupy a reproducible banana-shaped domain in the L*–b* plane, the physiologic basis for the confinement and boundaries of this region remains unknown. Second, although ITA assigns pigmentation categories from the angular position of a measured L*–b* coordinate, the causal relationship between this geometric construction and underlying melanin or blood content has not been established. In this work, we address both problems by establishing a physics-based mapping between skin chromophore parameters and CIELAB color coordinates. We simulate light–skin interactions using two complementary forward models parameterized by physiologically plausible melanin and blood concentrations, examine how isolated chromophore changes shift L*a*b* coordinates, and test whether the resulting geometry explains both ordinary skin-color variation and clinically relevant ITA failure modes.

We hypothesize that physiologically realistic variation in melanin and hemoglobin is sufficient to generate the observed banana-shaped distribution of human skin colors and to determine the directions and limits of color-coordinate shifts within CIELAB space. By independently varying these chromophores, the model defines iso-melanin and iso-blood trajectories, enabling quantification of perceptible color differences associated with blood variation across pigmentation levels. We further examine whether ITA remains specific for melanin along these trajectories and whether altered melanin depth can move coordinates outside the ordinary constitutive skin-color locus. More broadly, this physics-based mapping between chromophore parameters and colorimeter outputs provides a foundation for interpreting skin color measurements in terms of underlying physiology and for improving inference of melanin and hemoglobin from CIELAB coordinates.

## MATERIALS AND METHODS

### Generating reflectance curves spanning physiologic extremes

We adapted two previously published three-layer skin models for this study. The first used the adding-doubling (AD) method for photon transport (Else et al., 2024), and the second used MCmatlab, an open-source MATLAB implementation of a three-dimensional Monte Carlo (MC) algorithm based on Jacques’ MCXYZ solver (Marti et al., 2018; Jacques et al., 2017; Wang et al., 1995). Both models consisted of a melanosome-containing epidermis, blood-containing dermis, and hypodermis (Figure 1B). The computational geometry and layer-specific optical properties were kept identical across both models, and key model parameters are summarized in Table SI.

The epidermal melanosome volume fraction (mvf) was varied from 1% to 43%, and the dermal blood volume fraction (bvf) was varied from 0.2% to 7%. These ranges span physiologic extremes reported in the literature (Else et al., 2024; Jacques, 2013). We defined 10 logarithmically spaced levels within each range, yielding a 10 × 10 grid of (mvf, bvf) combinations (Figure 1B). For each combination, we ran both the AD and MC models with all other parameters held constant and computed the resulting diffuse reflectance spectrum at the skin surface. This procedure generated a database of 100 reflectance spectra per model, each linked to a specific (mvf, bvf) pair within the aforementioned range. This parameter sweep enabled isolation of chromophore-specific effects by examining one-dimensional slices through the grid. Specifically, varying bvf at fixed mvf defined iso-melanin trajectories, while varying mvf at fixed bvf defined iso-blood trajectories in CIELAB space.

Because multilayer skin models require assumptions about tissue geometry and optical properties, we performed sensitivity analyses to assess whether the predicted chromophore–color relationships were robust to plausible variations in these parameters. Specifically, we repeated the full mvf–bvf sweep across multiple layer-thickness combinations, including epidermal thickness of 60 and 120 µm and dermal thickness of 1, 2, and 3 mm (Table SI), spanning reported variation across anatomical sites (Oltulu, et al., 2018; Wang et al., 2025). We also evaluated sensitivity to tissue oxygenation by repeating the mvf–bvf sweep across oxygenation levels of 60%–80% using the AD model. These analyses were used to determine whether the key geometric features identified in this study reflect intrinsic chromophore interactions rather than artifacts of specific modeling assumptions.

While constitutive pigmentation is reasonably modeled by limiting melanin to the epidermis, several dermatologic conditions appear blue-gray because of substantial melanin or melanophages within the dermis. To evaluate whether the model could reproduce this optical regime, we performed an additional sensitivity analysis in which melanin absorption was introduced into the dermal layer while epidermal melanosome volume fraction and dermal blood volume fraction were varied as in the primary simulations. This analysis was designed to test whether altered melanin depth, rather than total melanin content alone, could shift the predicted skin-color locus outside the range generated by epidermal melanin and dermal blood.

### Extracting L*a*b* coordinates from reflectance curves

To map chromophore composition to perceived skin color, each simulated reflectance spectrum corresponding to a unique melanosome and blood volume fraction pair (mvf, bvf) was converted to CIELAB coordinates. Reflectance spectra were first transformed to XYZ tristimulus values using the CIE 10° standard observer color-matching functions and the relative spectral power distribution for the D65 illuminant (Kruschwitz, 2018). The XYZ values were then converted to L*a*b* coordinates, from which ITA values were calculated. This yielded a corresponding set of L*a*b* coordinates and ITA values for all 100 (mvf, bvf) combinations. The dermal melanin sensitivity simulations were processed using the same spectral-to-CIELAB pipeline, allowing the resulting coordinates and ITA values to be compared directly with the primary epidermal melanin simulations. We plotted these points in the L*-b* and L*-a* planes to visualize the simulated skin-color locus. For both the L*-a* and L*-b* planes, we compared the resulting distributions to results from the International Skin Spectra Archive (ISSA), a dataset comprised of spectral and colorimetric data from 2113 subjects across eight countries (Lu et al., 2025). The empirically defined ‘banana’ skin-color region from prior literature (Chardon et al., 1991; Del Bino at al., 2006) was also overlaid in the L*-b* plane to assess how well the model-generated locus aligned with reported skin-color boundaries. Then, for each model, we plotted the data individually to identify iso-blood and iso-melanin lines in each plane.

### Perceptible color differences (ΔE) from physiological variation in melanin and blood

To quantify perceptible color differences within the physiological range of each chromophore, ΔE was calculated while holding one chromophore constant and varying the other across its physiological limits. ΔE represents the Euclidean distance between two points in CIELAB color space (Kruschwitz, 2018):

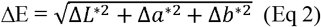

A ΔE value less than 2 is generally considered imperceptible to the human eye, while larger values correspond to increasingly noticeable color differences. To assess the perceptibility of blood-induced color changes (erythema) across different levels of pigmentation, ΔE was calculated between the minimum and maximum blood volume fractions (0.2% and 7%) while holding mvf fixed at representative values (1%, 5%, 10%, 20%, and 43%).

Conversely, to assess the perceptibility of pigmentation differences across varying blood content, ΔE was calculated between the minimum and maximum melanin volume fractions (1% and 43%) while holding bvf fixed at representative values (0.2%, 1%, 2%, 5%, and 7%).

### Evaluating ITA as a surrogate for melanin content

The use of ITA as an empirical surrogate for melanin content rests on an implicit geometric assumption: iso-melanin trajectories in the L*-b* plane are approximately linear and intersect at a common point of (*b*\*, *L*\*) = (0, 50). Under this assumption, points corresponding to a fixed melanin level (constant mvf) but varying blood volume (bvf) should lie along a straight line and yield similar ITA values.

Using the iso-melanin trajectories defined above, we evaluated this assumption in three ways. First, for each fixed mvf level (1%–43%), we assessed whether the corresponding trajectory in the *b*-L\** plane was well approximated by a straight line using linear regression and the Pearson correlation coefficient.

Second, we computed ITA at each point along these trajectories to quantify how ITA varies with bvf at a constant mvf. The spread of ITA values along each trajectory was used as a measure of blood-related confounding, with larger variation indicating greater sensitivity of ITA to changes in blood volume.

Finally, we examined whether allowing iso-melanin trajectories to intersect at a reference point (*b*_0_, *L*_0_) other than (0,50) could reduce blood-related confounding. ITA was therefore generalized to an adjusted form:

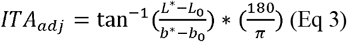

Inspection of the iso-melanin trajectories suggested that the portion where b* decreases with increasing bvf is approximately linear. We therefore restricted analysis to this regime and used linear regression and the Pearson correlation coefficient to quantify the degree of linearity across mvf levels. The intersection point (b□, L□) was estimated as the point of closest convergence among the resulting regression lines in the L*–b* plane.

Using this reference point, ITA_adj_ was computed for each point along the same iso-melanin trajectories, and the spread of ITA_adj_ values at fixed mvf as bvf varied and b* decreased was compared to that of the standard ITA to assess whether the adjusted formulation reduced blood-related confounding.

In addition to evaluating blood-related variation in ITA along iso-melanin trajectories, we separately assessed whether dermal melanin could create a distinct ITA failure mode by shifting simulated coordinates into the negative-b* region. Because standard ITA is calculated using b* in the denominator, measurements with negative b* may yield ITA values that reverse the expected relationship between increasing melanin and decreasing ITA. We therefore examined whether the dermal melanin sensitivity analysis produced negative b* values and whether these coordinates generated paradoxically high ITA values despite increased melanin absorption.

### Validation of model predictions

To evaluate whether experimentally observed color changes follow the model-predicted chromophore trajectories, we analyzed published datasets that reported CIE L*a*b* measurements under controlled physiological perturbations and assessed qualitative agreement between observed trajectories and model predictions. Data from Stamatas and Kollias was used to examine blood-driven color changes at approximately constant pigmentation (Stamatas and Kollias, 2004). In this study, changes in L*a*b* coordinates were reported from 10 subjects (Fitzpatrick skin types III–IV) during controlled venous occlusion at increasing cuff pressures, providing ΔL* and Δb* values across a range of blood volume states. Because absolute L* and b* coordinates were not directly reported, baseline values were estimated by combining the reported ΔL* and Δb* with independently reported ITA values at minimal and maximal occlusion. This enabled reconstruction of approximate iso-melanin trajectories in the L*–b* plane. Absolute a* values were not available; therefore, analysis in the L*–a* plane was limited to relative changes. All data were extracted by digitizing published figures.

Data from Park et al. were used to examine both blood-driven and melanin-driven color changes within the same cohort (Park et al., 1999). This study reported L*a*b* measurements from 30 subjects (Fitzpatrick skin types III–V) at baseline, 1 day after UV exposure, and 1 week after exposure. The baseline-to–day 1 measurements primarily reflect acute erythema due to increased blood volume with minimal change in melanin and were therefore used to approximate iso-melanin trajectories. In contrast, baseline-to–week 1 measurements reflect delayed tanning with increased melanin and resolution of erythema and were used to approximate iso-blood trajectories. Because subject-level coordinates were reported directly, trajectories in both the L*–b* and L*–a* planes were constructed without digitization or reconstruction.

In addition to published perturbation datasets, we evaluated two clinical examples representing predicted failure modes of ITA as a surrogate for melanin content. In each case, triplicate colorimetric measurements were obtained from lesional and adjacent unaffected skin using the Nix Spectro 2 Colorimeter, and median L*a*b* values were used for analysis. The first case involved a 67-year-old man with metastatic melanoma who presented with metastatic papules and multiple blue-black macules. Biopsy of one macule confirmed tumoral melanosis, a lesion characterized by abundant dermal melanin. This case was used to evaluate the model prediction that dermal melanin can shift b* toward negative values and produce paradoxical ITA values. The second case involved a 66-year-old woman with metastatic melanoma who presented with non-blanchable erythematous papules clinically diagnosed as leukocytoclastic vasculitis. This case was used to evaluate the complementary prediction that vascular-inflammatory color change can lower ITA despite no expected increase in melanin content.

## RESULTS

### Model-predicted skin color distributions in CIELAB space

To confirm physiologic plausibility of the AD and MC models, we examined layer-specific absorption and scattering properties together, with the resulting reflectance spectra across the mvf–bvf parameter space (Figures S1–S2). As expected from the model structure, epidermal absorption (µ_a_) varied only with melanin content, whereas dermal absorption varied only with blood volume. In the resulting reflectance spectra, increasing bvf primarily reduced overall reflectance while preserving spectral shape, whereas increasing mvf produced broadband attenuation that progressively suppressed hemoglobin-associated spectral features.

Having established that the models produce physiologically realistic reflectance spectra, we next examined how variations in melanin and blood volume mapped onto CIELAB color coordinates. Converting the simulated reflectance spectra across the generated 100-point mvf–bvf parameter space yielded a set of predicted L*a*b* coordinates spanning physiologic chromophore variation.

Figure 2 compares the distribution of model-predicted coordinates with empirical skin color measurements from the ISSA database. When plotted in the L*–b* plane (Figure 2A), both the AD and MC models generate a curvilinear region that closely matches the characteristic “banana-shaped” skin color volume locus first described by Chardon et al. and Del Bino et al. and subsequently confirmed by the ISSA dataset (Chardon et al., 1991; Del Bino et al., 2006; Lu et al., 2025). A similar correspondence is observed in the L*–a* plane (Figure 2B), where both models reproduce the structure of the empirical distribution. Despite differences in absolute positioning, the overall geometry is preserved across AD and MC models and aligns with the empirical distribution.

**Figure 2.**
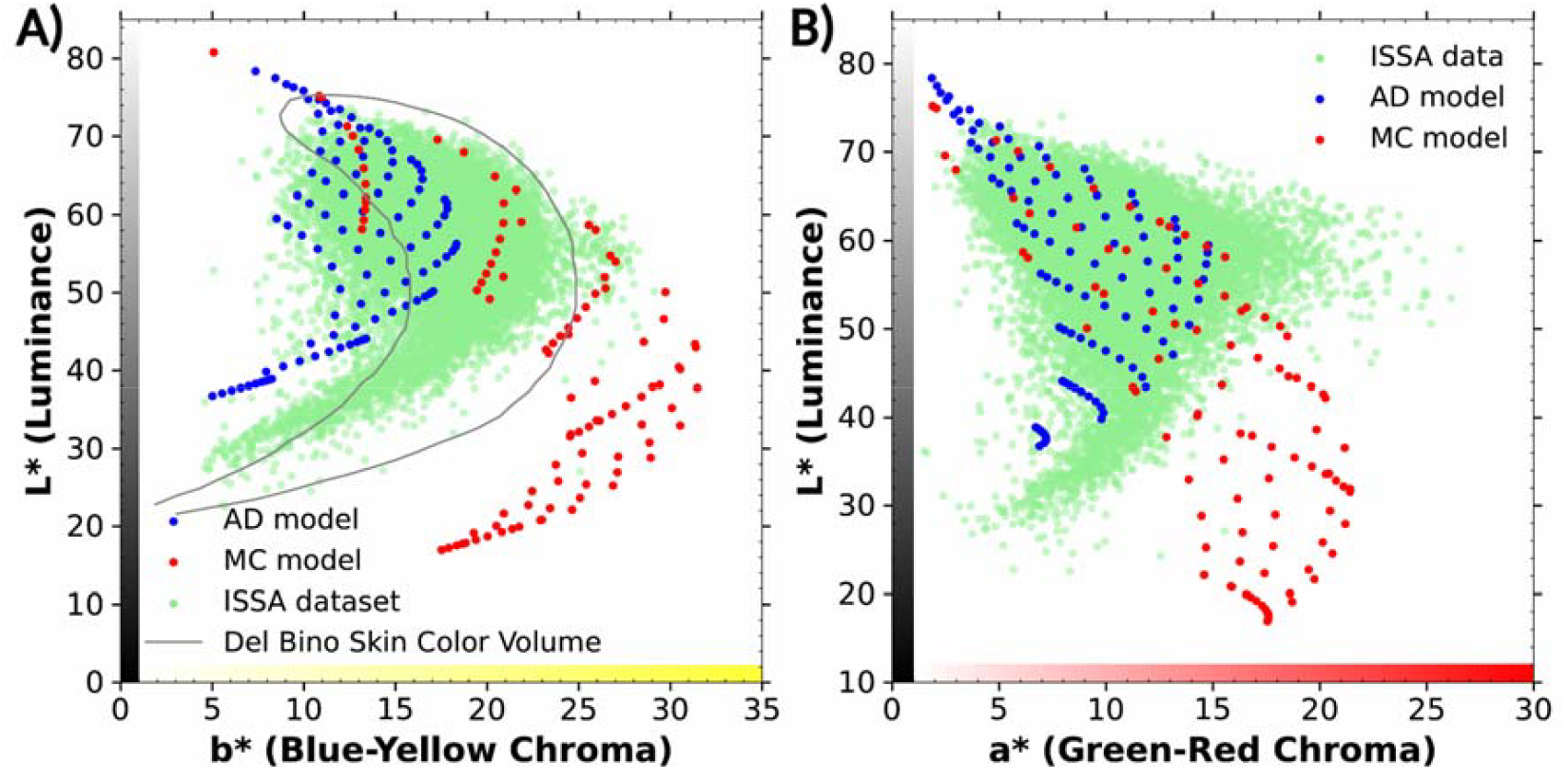
Model-predicted skin color distributions in CIELAB space compared with empirical colorimetry. Model-predicted L*a*b* coordinates were computed by converting simulated reflectance spectra for 100 combinations of melanosome volume fraction and blood volume fraction spanning physiologic ranges. Predicted coordinates from the adding–doubling (AD) model (blue) and Monte Carlo (MC) model (red) are overlaid on empirical skin colorimeter measurements from the ISSA database (green). (A) L*–b* plane. (B) L*–a* plane. In both projections, the distributions generated by the AD and MC models closely match the overall shape and extent of the empirical ISSA data. In the L*–b* plane, both models reproduce the characteristic curvilinear ‘banana-shaped’ locus of human skin colors (black curve in A): the distribution broadens and shifts toward higher b* as L* decreases from high to moderate values, then narrows and shifts toward lower b* as L* decreases further toward low values.

This agreement suggests that the empirically observed skin color locus arises from the optical interactions of melanin and hemoglobin and that variation in these chromophores across physiological extremes is sufficient to reproduce the characteristic region occupied by human skin in CIELAB space.

### Chromophore-specific trajectories in CIELAB space

We next examined chromophore-specific trajectories in the L*-b* plane. Figure 3A shows AD model-derived iso-melanin trajectories generated by varying bvf at fixed mvf (MC model results shown in Figure S3A). Figure 3B shows the complementary iso-blood trajectories generated by varying mvf at fixed bvf (MC results in Figure S3B). Across both trajectory types, the AD and MC models produce qualitatively similar patterns, with consistent geometric trends across modeling approaches, indicating that the observed trajectory geometry is not dependent on the specific optical simulation framework. For each iso-melanin trajectory, increasing bvf produces a decrease in L*, with the magnitude of this change decreasing as mvf increases, resulting in progressively compressed trajectories at higher melanin. In the b* dimension, iso-melanin trajectories are curved at lower mvf (1%, 5%, and 10%), with b* initially increasing and then decreasing as bvf increases, but become more linear at higher mvf (20%, 43%). On the other hand, each iso-blood trajectory appears to follow the overall curvature of the skin color locus, with a wider spread at low mvf and progressive convergence at higher mvf. Together, these results reveal distinct geometric signatures for melanin- and blood-driven variation in color space, providing a basis for separating their contributions to skin color coordinates.

**Figure 3.**
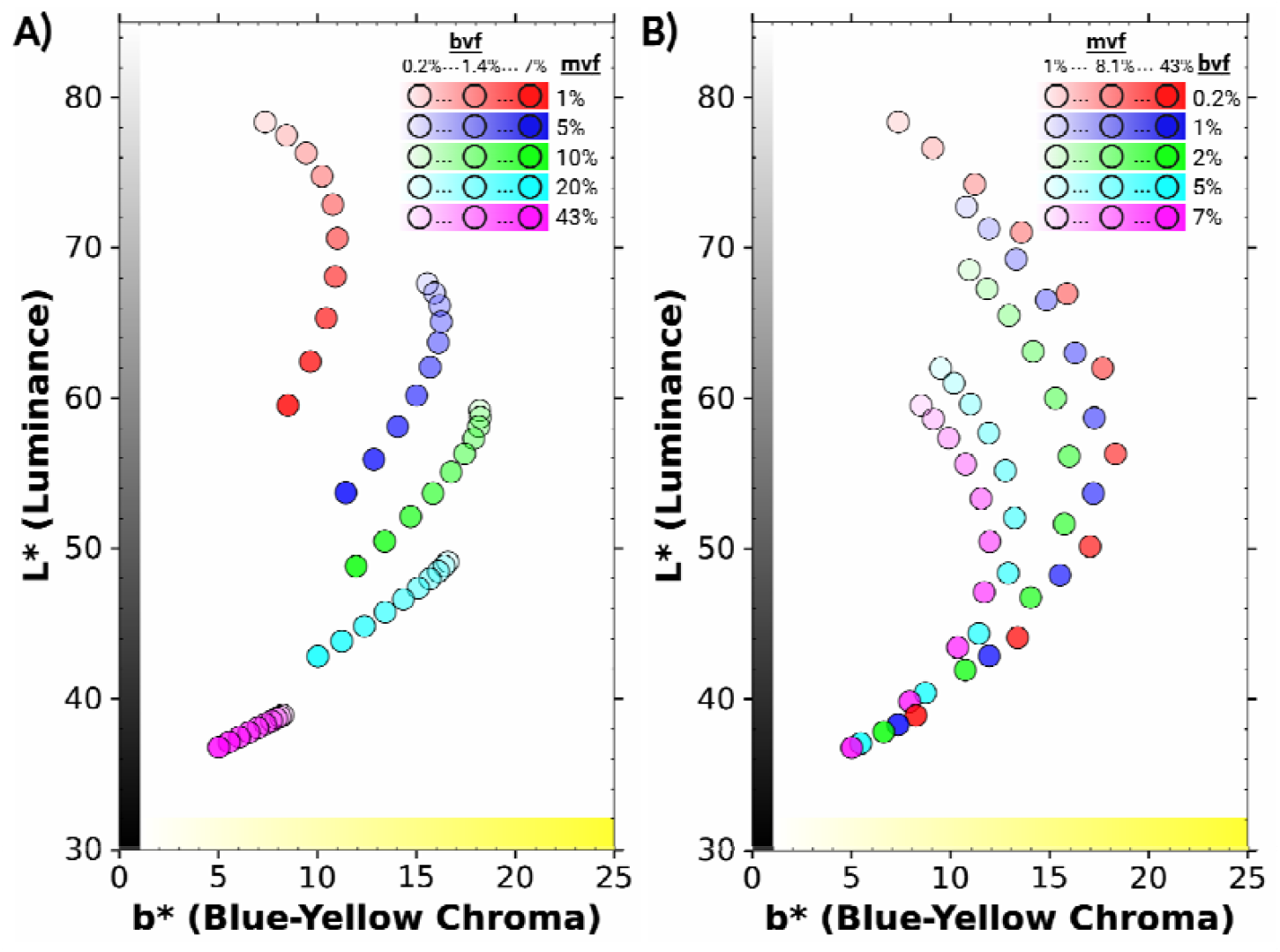
Iso-melanin and iso-blood trajectories in the L*b* plane. (A) Iso-melanin trajectories in the L*–b* plane generated by holding melanosome volume fraction (mvf) constant while varying blood volume fraction (bvf) using the adding–doubling (AD) model. Colored points denote fixed mvf levels with increasing bvf. Increasing opacity of the colored points indicates increasing bvf for a given mvf level. (B) Iso-blood trajectories generated by holding bvf constant while varying mvf using the AD model. Colored points denote fixed bvf levels with increasing mvf. Increasing opacity of the colored points indicates increasing mvf for a given bvf level. Results are shown for the adding–doubling (AD) model; corresponding Monte Carlo results are provided in Figure S3.

Similar geometric relationships were observed in the L*–a* plane (Figure 4). Iso-melanin trajectories generated by varying bvf at fixed mvf showed a wide spread in a* at low mvf that progressively compressed at higher mvf, indicating reduced sensitivity of a* to blood variation with increasing melanin. Iso-blood trajectories generated by varying mvf at fixed bvf demonstrated steeper decline in L* with increasing a* compared to iso-melanin trajectories As mvf increased, iso-blood trajectories converged toward a narrower region of color space. As in the L*–b* plane, the overall coordinate spread was largest at low mvf (or low bvf) and progressively decreased at higher mvf, reflecting suppression of blood-driven color variation with increasing melanin. Comparable patterns were observed in the MC model (Figure S4), indicating that these geometric relationships are robust to the modeling approach.

**Figure 4.**
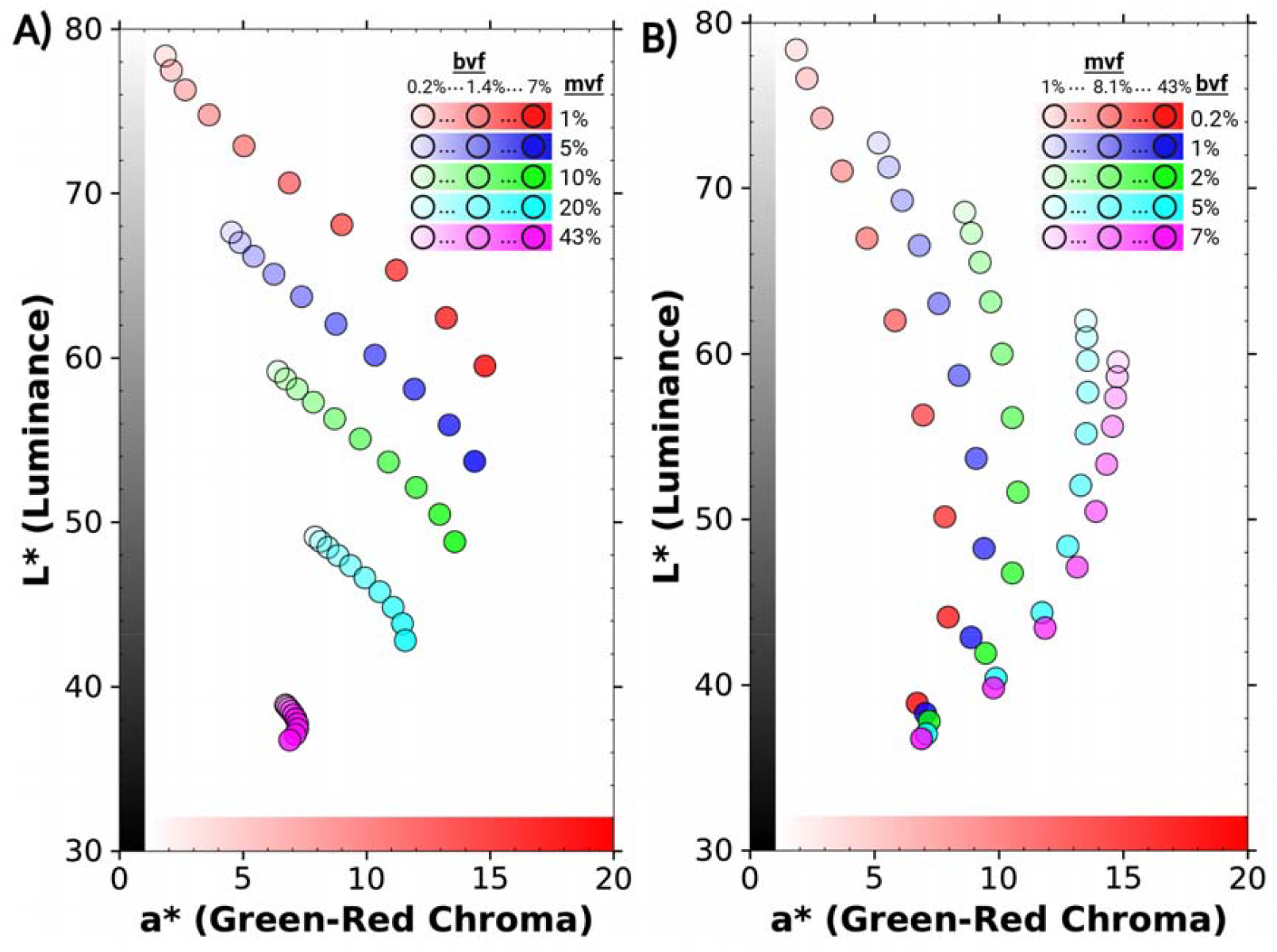
Iso-melanin and iso-blood trajectories in the L*–a* plane. (A) Iso-melanin trajectories obtained by varying blood volume fraction (bvf) at fixed melanosome volume fraction (mvf). Each colored curve represents a constant mvf level across increasing bvf. Increasing opacity of the colored points indicates increasing bvf for a given mvf level. (B) Iso-blood trajectories obtained by varying mvf at fixed bvf. Each colored curve represents a constant bvf level across increasing mvf. Increasing opacity of the colored points indicates increasing mvf for a given bvf level. Results are shown for the adding–doubling (AD) model; corresponding Monte Carlo results are provided in Figure S4.

### Perceptible color differences along iso-melanin and iso-blood trajectories

To relate these geometric patterns to visual perception, we quantified color differences (ΔE) computed from AD model–derived CIELAB coordinates across the physiological ranges of melanin (mvf) and blood volume (bvf) (Figure 5). When comparing the extremes of melanin content (1% vs 43% mvf) at a range of fixed bvf values (Figure 5A), ΔE remained large across the full range of blood volumes, decreasing from 39.8 at 0.2% bvf to 24.3 at 7% bvf. Thus, variation in melanin produces strongly perceptible color differences regardless of blood content. In contrast, when comparing the extremes of blood volume (0.2% vs 7% bvf) at a range of fixed mvf values (Figure 5B), ΔE decreased markedly with increasing melanin, falling from 22.8 at 1% mvf to 3.8 at 43% mvf. Similar trends were observed with the MC model (Figure S5). These results show that increasing melanin progressively suppresses the perceptible color change associated with varying blood volume, whereas varying melanin produces large perceptible color differences across the full range of blood volumes.

**Figure 5.**
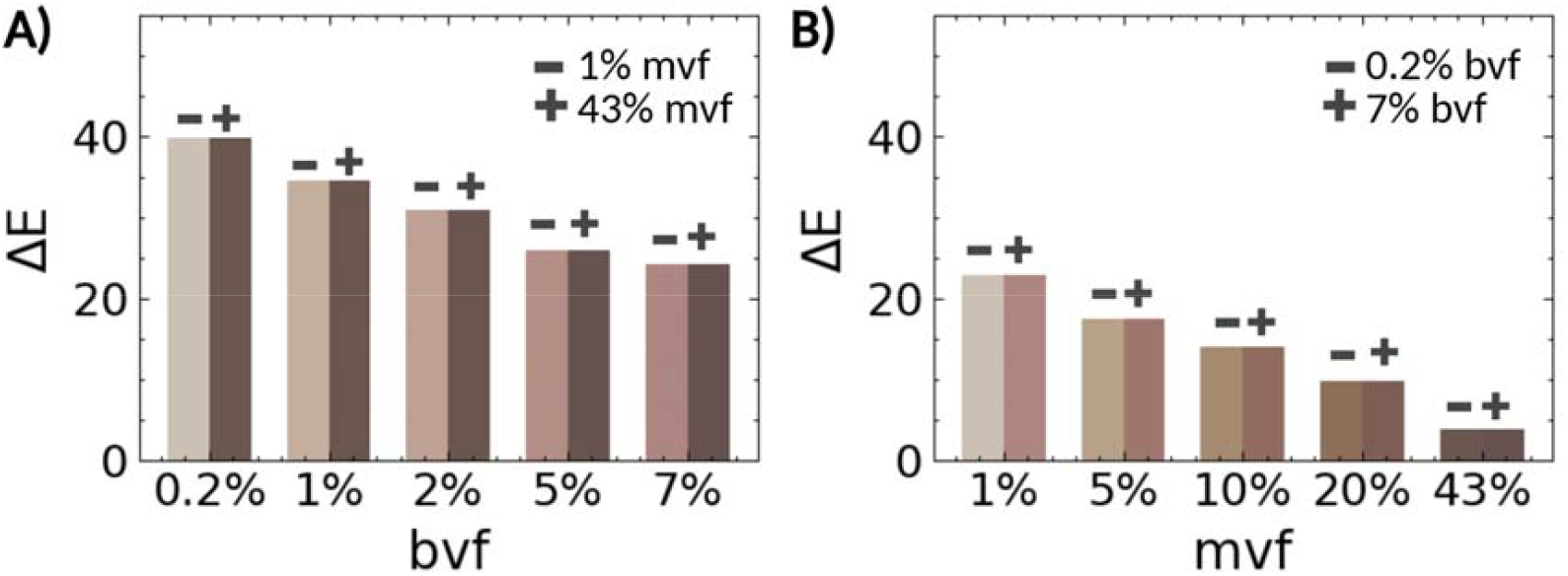
Perceptible color differences (ΔE) produced by physiological variation in blood and melanin. (A) ΔE between the minimum and maximum physiological melanosome volume fraction (mvf: 1-43%) at fixed blood volume fraction (bvf) levels predicted by the adding–doubling (AD) model. (B) ΔE between the minimum and maximum physiological blood volume fraction (bvf: 0.2–7%) at fixed mvf levels predicted by the AD model. Each bar represents the color difference between the L*a*b* coordinates corresponding to the minimum and maximum chromophore values used in the comparison; the two colors indicate the respective L*a*b* values of those endpoints. ΔE associated with variation in blood volume decreases as mvf increases, whereas ΔE associated with variation in melanin remains large across the full range of bvf. Corresponding results for the MC model in Figure S5.

### Exploring the relationship between melanin and ITA

For ITA to serve as a surrogate for melanin, it should remain constant along iso-melanin trajectories, which, based on Equation 2, requires iso-melanin trajectories in the L*–b* plane to be linear and intersect at (*b*\*, *L*\*) = (0, 50).

We evaluated this requirement using the simulated iso-melanin trajectories. Linear regression of full iso-melanin trajectories showed that goodness of fit increased markedly with mvf. For the AD model, R^2^ values increased from 0.05 at 1% mvf to 0.83 at 5% mvf, and to ≥0.97 at mvf ≥10% (Figure 6A), with similar trends observed in the MC model (Figure S6). These results indicate that iso-melanin trajectories lack linearity at low mvf but become nearly linear at higher mvf. Consistent with this, substantial variation in ITA was observed along iso-melanin trajectories at low mvf (Figure 6B–C). For example, at 1% mvf, ITA ranged from 48° to 76° (ΔITA = 28°), and at 5% mvf, from 18° to 49° (ΔITA = 31°). This variation decreased with increasing mvf but remained non-negligible even at high melanin levels (ΔITA = 16° at 43% mvf), indicating that standard ITA is sensitive to blood volume, particularly at low mvf.

**Figure 6.**
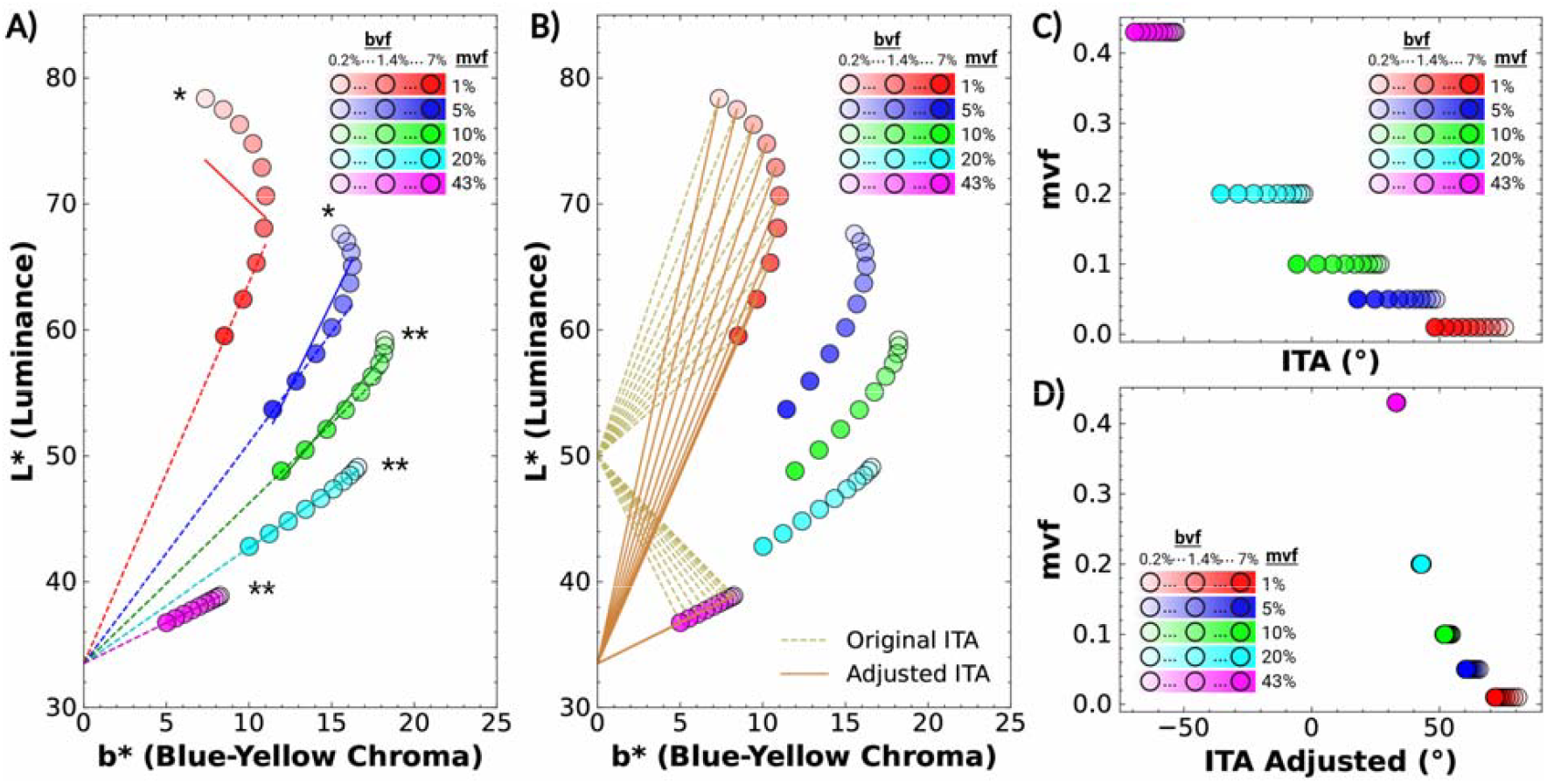
Evaluation of ITA as a surrogate for melanin. (A) Linear regression of iso-melanin trajectories in the L*–b* plane for the AD model. Colored points represent simulated L*–b* coordinates along iso-melanin trajectories generated by varying blood volume fraction (bvf) at fixed melanosome volume fraction (mvf). Increasing opacity of the colored points indicates increasing bvf for a given mvf. Solid lines show linear fits across the full trajectories (* R^2^<0.97, ** R^2^≥ 0.97), while dashed lines show fits restricted to the portion of each trajectory where b* decreases with increasing bvf. Restricting the regression to this region improves linearity and produces convergence of the fitted lines toward a common intersection point. (B) Geometric interpretation of standard ITA and adjusted ITA. Lines corresponding to constant ITA values are drawn from the standard intercept at (b*, L*) = (0, 50) (dashed lines) and from the empirically determined intersection point (b□, L□) = (0, 33.5) (solid lines). Examples are shown for the 1% and 43% iso-melanin trajectories. (C) ITA values calculated using the standard formulation (Eq. 1) for all simulated points along iso-melanin trajectories. Substantial variation in ITA occurs along each iso-melanin trajectory, particularly at low mvf levels. (D) ITA values calculated using the adjusted formulation (Eq. 3). The spread of ITA values along iso-melanin trajectories is substantially reduced, particularly at higher mvf levels.

When analysis was restricted to the regime where *b*\* decreases with increasing bvf (dotted lines Figure 6A), linearity improved across all iso-melanin trajectory levels (R^2^ ≥ 0.97). Regression lines from this regime converged toward a common region in the L*–b* plane, yielding an approximate intersection at (*b*□, *L*□) = (0, 33.5) for the AD model, with a similar convergence observed in the MC model (Figure S6).

Together, these results show that the geometric assumptions underlying ITA are not satisfied across the full chromophore range, resulting in substantial blood-driven variation in ITA, particularly at low melanin levels, although this effect is reduced as trajectories become more linear at higher mvf. To reduce blood-related variation, we defined a generalized ITA formulation (ITA_adj_) by replacing the standard reference point (0, 50) with the empirically determined intersection point (b□, L□) = (0, 33.5), consistent with Equation 3 (Figure 6B). Across the full chromophore space, the total dynamic range of ITA_adj_ was smaller than that of standard ITA (48° vs 144°), reflecting compression of the metric scale. As a result, although the absolute spread of ITA values along iso-melanin trajectories decreased (Figure 6D), this reduction was less pronounced when considered relative to the total dynamic range, particularly at low mvf where trajectories are curved. For example, at 1% mvf, variation remained ∼19% of the total dynamic range for both standard ITA and ITA_adj_. In contrast, when analysis was restricted to the regime where *b*\* decreases with increasing bvf, ITA_adj_ substantially reduced variation across all iso-melanin trajectories. In this regime, the relative variation (ΔITA / total range) for ITA_adj_ versus standard ITA was 2.1% vs 7.6% at 1% mvf, 0.2% vs 11.1% at 5% mvf, 0.4% vs 13.2% at 10% mvf, 1.0% vs 16.0% at 20% mvf, and 0.02% vs 6.3% at 43% mvf.

These findings show that variability of ITA arises from a geometric mismatch between the assumed intercept of the standard formulation and the actual structure of iso-melanin trajectories. Adjusting the reference point improves melanin specificity in the regime where the trajectories are linear, but it cannot fully correct for variation at low melanin levels, where the trajectories are intrinsically curved.

To evaluate whether these geometric relationships were robust to modeling assumptions, we performed sensitivity analyses varying epidermal thickness, dermal thickness, and tissue oxygenation across physiologically plausible ranges (Supplementary Figures S7–S10). Across all perturbations, the dominant geometric features of the simulated color space—including the structure of iso-melanin and iso-blood trajectories and the linearity of high-melanin iso-melanin trajectories—were preserved, indicating that the chromophore–color relationships identified here are robust to these model parameters.

We also evaluated a distinct ITA failure mode predicted by altering melanin depth. When melanin absorption was introduced into the dermal layer, the simulated L*–b* trajectories extended beyond the ordinary constitutive skin-color locus and crossed into negative b* values, particularly when epidermal melanosome volume fraction was low (Figure S11). Because standard ITA uses b* in the denominator, negative b* values reversed the expected directionality of the metric: instead of producing lower ITA values with increasing tissue melanin, dermal melanin could yield paradoxically high positive ITA values. Thus, the model predicts that melanin-rich blue-gray lesions may be misclassified by standard ITA as having low melanin content.

### External and clinical validation of model predictions

Stamatas and Kollias reported the mean L* and b* values from 10 subjects measured at the forearm during upper arm blood pressure cuff occlusion, ranging from 0 to 60 mmHg (Stamatas and Kollias, 2004). Because venous occlusion changes blood content while melanin remains constant, these data provide a real-world iso-melanin trajectory. When reconstructed in the L*–b* plane, the trajectory agreed with the model-predicted iso-melanin geometry and was highly linear (R^2^ = 0.99; Figure 7A). Notably, the trajectory did not intersect the standard ITA reference point at (b*, L*) = (0, 50), supporting the conclusion that blood-driven shifts can alter ITA despite unchanged melanin.

**Figure 7.**
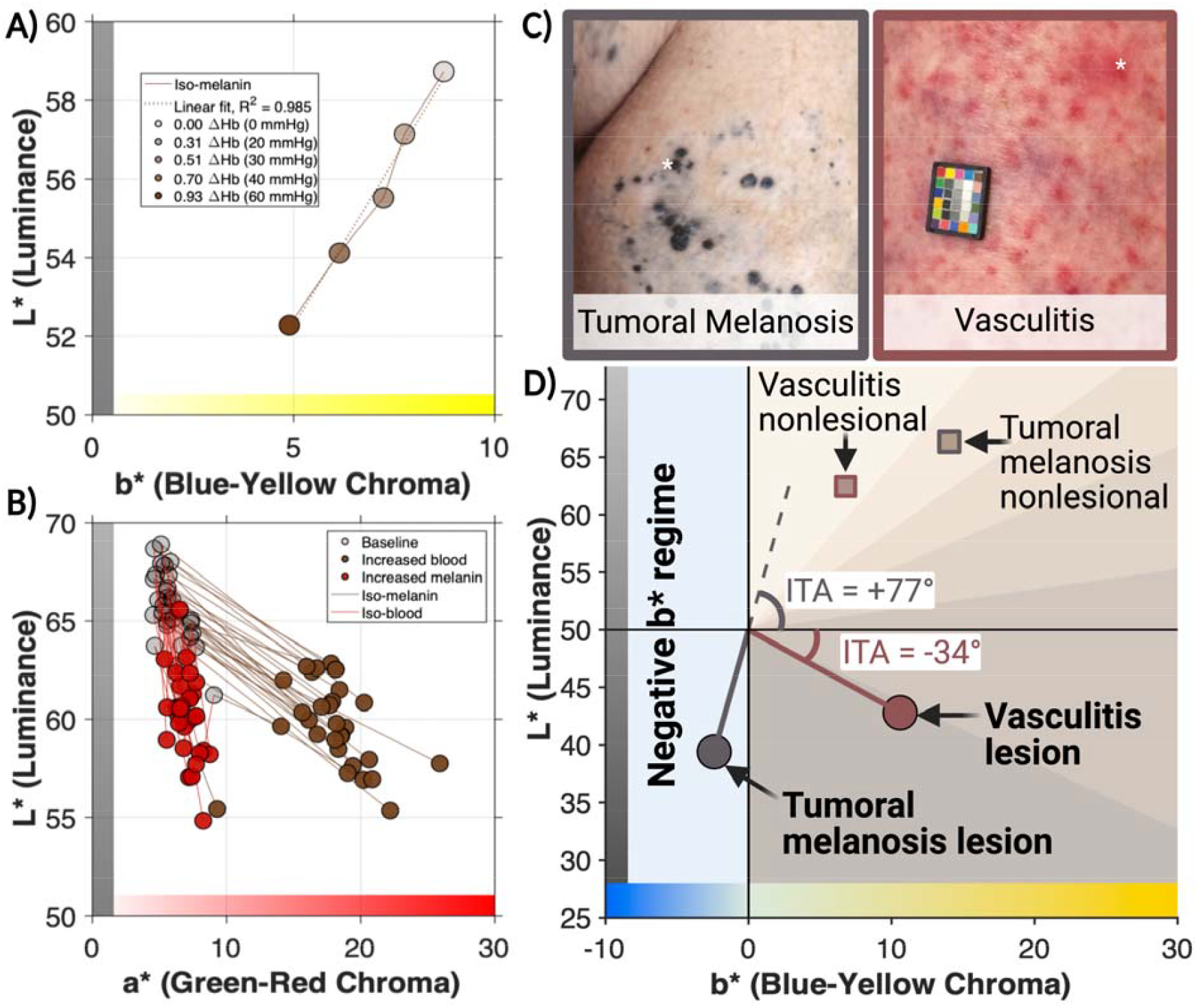
Validation of model-predicted skin color trajectories and ITA failure modes. (A) Reconstructed iso-melanin trajectory in the L*–b* plane from Stamatas and Kollias during blood-pressure-cuff-induced venous congestion. Because melanin remains constant while blood content changes, the observed trajectory provides an empirical test of the model-predicted iso-melanin direction. The trajectory falls within the regime where b* decreases with decreasing L* and is highly linear. (B) Baseline-to-day-1 shifts reflect acute erythema with minimal expected melanin change and therefore approximate iso-melanin trajectories, whereas baseline-to-week-1 shifts reflect delayed pigmentation change after resolution of acute erythema and therefore approximate iso-blood trajectories. The shallower iso-melanin and steeper iso-blood directions are consistent with model-predicted chromophore-specific geometry. (C) Representative clinical photographs of tumoral melanosis and vasculitis lesions. White markers indicate approximate colorimetry sites. (D) Clinical colorimetry demonstrates two predicted ITA failure modes. The vasculitis lesion shifted from very light constitutive pigmentation to a dark-category ITA despite no expected increase in melanin, consistent with blood-related confounding. The tumoral melanosis lesion shifted into the negative-b* region and yielded a paradoxically high ITA despite abundant dermal melanin.

The Park et al. dataset provided complementary validation in the L*–a* plane (Park et al., 1999). This study reported colorimeter-derived L*a*b* values from 30 subjects at baseline, 1 day after UV exposure, and 1 week after UV exposure. Baseline-to-day-1 shifts, interpreted as acute erythema with minimal melanin change, approximated iso-melanin trajectories, whereas baseline-to-week-1 shifts, interpreted as tanning-related delayed pigmentation after resolution of acute erythema, approximated iso-blood trajectories. Consistent with the model predictions, the iso-melanin trajectories were shallower than the iso-blood trajectories in the L*–a* plane (Figure 7B).

Finally, the two clinical cases demonstrated predicted ITA failure modes in dermatologic lesions (Figure 7C-D). In the vasculitis case, nonlesional skin had median L*a*b* values of (62.4, 9.6, 6.8), corresponding to an ITA of +61° and a very light constitutive pigmentation category. In contrast, the vasculitis papule had median L*a*b* values of (42.8, 26.3, 10.6), corresponding to an ITA of −34° and a dark pigmentation category. Because this was a vascular-inflammatory lesion with no expected increase in melanin content, the low lesional ITA reflects blood-related confounding rather than increased melanin.

In the tumoral melanosis case, nonlesional skin had median L*a*b* values of (66.4, 4.8, 14.0), corresponding to an ITA of +49° and a light constitutive pigmentation category. In contrast, the tumoral melanosis lesion had median L*a*b* values of (39.4, 4.1, −2.4). Because tumoral melanosis contains abundant dermal melanin, the lesion would be expected to yield a low ITA if ITA remained a faithful surrogate for melanin content. Instead, the negative lesional b* value produced a calculated ITA of +77°, a value ordinarily associated with very light pigmentation and low melanin content. Thus, the clinical cases demonstrate that vascular-inflammatory color change can artifactually decrease ITA, whereas dermal melanin can artifactually increase ITA when b* becomes negative.

## DISCUSSION

In this study, we developed a physics-based mapping between skin chromophore content and CIELAB coordinates that provides a mechanistic explanation for two long-standing observations in skin colorimetry. First, physiologically plausible variation in melanin and blood volume was sufficient to reproduce the characteristic ‘banana-shaped’ locus of human skin colors in the L*–b* plane. Second, the same framework revealed that melanin and blood generate distinct trajectories in color space, allowing chromophore-specific contributions to skin color to be interpreted geometrically. Together, these findings move the interpretation of colorimeter outputs beyond empirical correlation and towards a causal optical framework grounded in skin physiology.

A key strength of this framework is that its predictions were preserved across two distinct light-transport approaches. Although the adding–doubling and Monte Carlo models differed in the absolute position of the predicted loci, they showed strong agreement in the features most relevant to interpretation: the overall skin-color locus, the curvilinear nature of iso-blood trajectories, the increasing linearity of iso-melanin trajectories at high melanin, and the compression of blood-driven color variation as melanin increases. Sensitivity analyses further demonstrated that these geometric relationships were stable across plausible variations in epidermal thickness, dermal thickness, and tissue oxygenation. Thus, while absolute color coordinates depend on modeling assumptions, the trajectory geometry identified here appears to be a robust consequence of the optical interaction between melanin and hemoglobin.

These findings are consistent with prior empirical observations of chromophore-specific color changes under physiological perturbation. In the present study, we reanalyzed published data from Stamatas and Kollias and from Park et al. and found that the observed color changes follow the chromophore-specific trajectory geometry predicted by our model (Stamatas and Kollias, 2004; Park et al., 1999). Independent studies report similar behavior. For example, Takiwaki et al. observed that both iso-melanin and iso-blood trajectories are approximately linear in the L*–a* plane, with iso-melanin trajectories exhibiting a shallower slope than iso-blood trajectories, in agreement with our theoretical predictions (Takiwaki et al., 2002). Together, these observations support the generality of the chromophore-specific trajectory geometry identified here.

The two clinical cases extend this validation to dermatologic lesions. The vasculitis case paralleled the venous-congestion data from Stamatas and Kollias: vascular-inflammatory color change lowered ITA into a range ordinarily interpreted as high melanin content despite no expected increase in melanin. Conversely, the tumoral melanosis case reproduced the distinct failure mode predicted by the dermal melanin sensitivity analysis. Introducing melanin absorption into the dermis extended the L*–b* trajectory into negative b* values and produced paradoxically high positive ITA values despite increased melanin absorption. Similarly, the tumoral melanosis lesion had a negative b* value consistent with blue-gray dermal melanin, yet its calculated ITA fell in a range ordinarily associated with very light pigmentation. These cases show that ITA can fail in opposite directions depending on the underlying chromophore change: blood or inflammation can artifactually decrease ITA, whereas dermal melanin can artifactually increase ITA when b* becomes negative.

Our results clarify both the strengths and limitations of the Individual Typology Angle as a surrogate for melanin content. Standard ITA assumes that iso-melanin trajectories are linear and intersect at a fixed point in the L*–b* plane. Our simulations show that this approximation has a physiological basis at higher melanin levels, where iso-melanin trajectories become strongly linear, but it breaks down at lower melanin levels, where trajectories are curved, and the apparent intersection does not occur at b* = 0, L* = 50. As a result, standard ITA can vary substantially with blood volume even when melanin is constant. By shifting the ITA reference point toward the empirically determined intersection of the approximately linear high-melanin trajectories, we substantially reduced blood-driven variation relative to the total dynamic range of the metric. This suggests that ITA is best understood not as a fixed empirical construct but as a geometric measure whose melanin specificity depends on how well its reference geometry matches the underlying chromophore trajectories.

The dermal melanin analysis further shows that ITA depends on whether the measured coordinate remains within the ordinary positive-b* constitutive skin-color locus. When b* becomes negative, as may occur in blue-gray dermal pigmentary lesions, the standard ITA formula can reverse the expected relationship between melanin and ITA. Replacing b* with |b*| would leave ITA unchanged for ordinary positive-b* measurements while preventing this sign-reversal failure mode when b* is negative. Thus, an absolute-b* formulation may provide a simple and conservative modification that preserves the expected directionality between melanin-rich blue-gray pigmentation and lower ITA-like values. However, this approach should still be validated across lesion types and devices before being adopted as a general replacement for standard ITA.

Several limitations should be acknowledged. Although the two optical models agreed strongly in their geometric predictions, both simplify the complexity of real skin and do not capture all biological sources of variation. Blood oxygenation was varied only within the adding–doubling framework, although the modest effects observed and the strong agreement between the two models suggest that this does not alter the primary geometric relationships identified here. The dermal melanin analysis was also simplified. It was intended to test whether altered pigment depth could shift the colorimetric locus into the negative-b* region, not to fully model the optical complexity of lesions characterized by abundant dermal melanin. Therefore, the dermal melanin simulations should be interpreted as a sensitivity analysis demonstrating a plausible geometric failure mode of ITA rather than a complete model of tumoral melanosis or other blue-gray lesions. The clinical cases should likewise be interpreted as illustrative examples rather than a validation cohort. They demonstrate that the predicted ITA failure modes can occur in real dermatologic lesions, but larger studies are needed to determine how often these errors occur across lesion types, pigmentation backgrounds, anatomic sites, and devices. The adjusted ITA formulation also involves a tradeoff: shifting the reference point reduced blood-driven variation along iso-melanin trajectories, but compressed the overall ITA dynamic range, which may increase sensitivity to measurement noise or small coordinate errors.

Finally, the ΔE analysis reflects perceptible color differences produced by physiological variation in dermal blood volume within the modeled range of 0.2–7% bvf. These bounds do not necessarily encompass the full range of blood content that may occur in pathological states such as inflammation, vascular malformations, hemorrhage, or venous congestion. Future work extending the analysis to higher blood volume fractions and more complex vascular geometries may yield larger absolute ΔE values. However, because the suppression of blood-driven color variation arises from broadband attenuation produced by epidermal melanin, the relative reduction in perceptible color change with increasing pigmentation would be expected to persist even at higher blood volumes. Accordingly, the qualitative conclusion that erythema becomes progressively less perceptible as melanin increases is robust, although the absolute magnitude of ΔE differences should be interpreted within the limits of the modeled physiological ranges.

In summary, this work establishes a causal optical framework linking melanin and blood concentrations to CIELAB skin color coordinates. By explaining the origin of the ‘banana-shaped’ skin-color locus, defining the geometry of chromophore-specific trajectories, and showing how blood-driven color differences become compressed at high melanin, our results unify several previously empirical observations within a single mechanistic model of skin color. This framework clarifies why erythema is more difficult to detect in highly pigmented skin, why ITA is confounded by blood volume, and why ITA can fail when dermatologic lesions alter pigment depth or move b* outside the ordinary constitutive skin-color range. More broadly, these findings suggest that skin colorimetry can move beyond descriptive classification toward quantitative inference of underlying chromophore physiology. Such an approach may enable more interpretable optical biomarkers, more equitable diagnostic tools, and improved device performance across the full spectrum of human skin pigmentation.

## CONFLICT OF INTEREST

The authors have no conflicts of interest to disclose.

## ACKNOWLEDGEMENTS

This work was supported by funding from the National Institutes of Health DP5OD031872 and by institutional support from Washington University in St. Louis.

## AUTHOR CONTRIBUTIONS

Conceptualization: M.H., Y.H., M.S., Z.N., L.S.; Data Curation: M.H, Y.H.; Formal Analysis: M.H., Y.H., M.S., Z.N., L.S.; Funding Acquisition: L.S.; Investigation: M.H., Y.H.; Methodology: M.H., Y.H., M.S., Z.N., L.S.; Project Administration: L.S.; Resources: Z.N., L.S.; Software: M.H., Y.H., M.S.; Supervision: L.S. Visualization: M.H., Y.H., L.S.; Writing – original draft: M.H., Y.H., L.S.; Writing – review & editing: M.H., Y.H., M.S., Z.N., L.S.

## Notes

### Competing Interest Statement

The authors have declared no competing interest.

## REFERENCES

Alaluf S, Atkins D, Barrett K, Blount M, Carter N, Heath A. The Impact of Epidermal Melanin on Objective Measurements of Human Skin Colour. Pigment Cell Research. 2002 Apr;15(2):119–26. doi:10.1034/j.1600-0749.2002.1o072.x

Chardon A, Cretois I, Hourseau C. Skin colour typology and suntanning pathways. Intern J of Cosmetic Sci. 1991 Aug;13(4):191–208. doi:10.1111/j.1467-2494.1991.tb00561.x

Del Bino S, Bernerd F. Variations in skin colour and the biological consequences of ultraviolet radiation exposure. British Journal of Dermatology. 2013;169(3):33–40. doi:10.1111/bjd.12529

Del Bino S, Sok J, Bessac E, Bernerd F. Relationship between skin response to ultraviolet exposure and skin color type. Pigment Cell Research. 2006 Dec;19(6):606–14. doi:10.1111/j.1600-0749.2006.00338.x

Else TR, Hacker L, Gröhl J, Bunce EV, Tao R, Bohndiek SE. Effects of skin tone on photoacoustic imaging and oximetry. J Biomed Opt. 2024 Jan;29(Suppl 1):S11506. doi:10.1117/1.JBO.29.S1.S11506 PubMed PMID: 38125716; PubMed Central PMCID: PMC10732256.

Feiner JR, Severinghaus JW, Bickler PE. Dark Skin Decreases the Accuracy of Pulse Oximeters at Low Oxygen Saturation: The Effects of Oximeter Probe Type and Gender. Anesthesia & Analgesia. 2007 Dec;105(6):S18–23. doi:10.1213/01.ane.0000285988.35174.d9

Hou S, Li A. Skin color of Chinese women across different regions of China: An analysis based on both individual typology angle and hue angle. Journal of Dermatologic Science and Cosmetic Technology. 2024 Mar;1(1):100003. doi:10.1016/j.jdsct.2024.100003

Jacques SL. Optical properties of biological tissues: a review. Phys Med Biol. 2013 Jun 7;58(11):R37–61. doi:10.1088/0031-9155/58/11/R37

Jacques SL, Li T, Prahl, S. mcxyz.c, a 3D Monte Carlo simulation of heterogeneous tissues. 2017. http://omlc.org/software/mc/mcxyz.

Kruschwitz JD. Field Guide to Colorimetry and Fundamental Color Modeling. SPIE; 2018 doi:10.1117/3.2500912

Ly BCK, Dyer EB, Feig JL, Chien AL, Del Bino S. Research Techniques Made Simple: Cutaneous Colorimetry: A Reliable Technique for Objective Skin Color Measurement. Journal of Investigative Dermatology. 2020 Jan;140(1):3–12.e1. doi:10.1016/j.jid.2019.11.003

Lu Y, Xiao K, Pointer M, He R, Zhou S, Nasseraldin A, et al. The International Skin Spectra Archive (ISSA): a multicultural human skin phenotype and colour spectra collection. Sci Data. 2025 Mar 23;12(1):487. doi:10.1038/s41597-025-04857-5

Marti D, Aasbjerg RN, Andersen PE, Hansen AK. MCmatlab: an open-source, user-friendly, MATLAB-integrated three-dimensional Monte Carlo light transport solver with heat diffusion and tissue damage. J Biomed Opt. 2018 Dec 15;23(12):1. doi:10.1117/1.JBO.23.12.121622

Oltulu P, Ince B, Kokbudak N, Findik S, Kilinc F. Measurement of epidermis, dermis, and total skin thicknesses from six different body regions with a new ethical histometric technique. Turk J Plast Surg. 2018;26(2):56. doi:10.4103/tjps.TJPS_2_17

Park SB, Suh DH, Youn JI. A long-term time course of colorimetric evaluation of ultraviolet light-induced skin reactions: Colorimetric evaluation of UV-induced skin reactions. Clinical and Experimental Dermatology. 1999 Jul;24(4):315–20. doi:10.1046/j.1365-2230.1999.00488.x

Pryor Y, He J, Kang J, Jenkins B, Lasisi T. Optical Limits in Skin Reflectance Measurement: Quantifying Melanin-Dependent Constraints on Erythema Detection. bioRxiv. 2025 Dec. doi: 10.64898/2025.12.22.696093

Stamatas GN, Kollias N. Blood stasis contributions to the perception of skin pigmentation. J Biomed Opt. 2004;9(2):315. doi:10.1117/1.1647545

Takiwaki H, Miyaoka Y, Kohno H, Arase S. Graphic analysis of the relationship between skin colour change and variations in the amounts of melanin and haemoglobin. Skin Research and Technology. 2002 May;8(2):78–83. doi:10.1034/j.1600-0846.2002.00333.x

Wang LV, Jacques SL, Zheng L. MCML – Monte Carlo modeling of light transport in multi-layered tissues. Comput. Methods Prog. Biomed. 1995 Apr; 47: 131–146. doi: 10.1016/0169-2607(95)01640-F

Wang Y, Tian M, Guo R, Du F, Qiu L, Tang Y. Quantification of normal skin thickness using very high-frequency ultrasound: a clinical study in Chinese adults. Quant Imaging Med Surg. 2025 Jun;15(6):5218–31. doi:10.21037/qims-2024-2637

Zonios G, Bykowski J, Kollias N. Skin Melanin, Hemoglobin, and Light Scattering Properties can be Quantitatively Assessed In Vivo Using Diffuse Reflectance Spectroscopy. Journal of Investigative Dermatology. 2001 Dec 1;117(6):1452–7. doi:10.1046/j.0022-202x.2001.01577.x

