## Supplemental Materials for "The optical origin of the human skin color ‘banana’ in CIELAB space"

Table SI. Parameters used in both models.

| **Parameter** | **Value** |
| --- | --- |
| Epidermal thickness | 60 µm, 120µm |
| Dermal thickness | 1 mm, 2mm, 3mm |
| Hypodermis thickness | Infinite |
| Scattering anisotropy epidermis | 0.9 |
| Scattering anisotropy dermis | 0.9 |
| Scattering anisotropy hypodermis | 0.7 |
| Dermal water volume fraction | 65% |
| Hypodermis water volume fraction | 65% |
| Melanosome volume fraction range (epidermis) | [1, 1.5, 2.3, 3.5, 5.3, 8.1, 12.3, 18.6, 28.3, 43%] |
| Blood volume fraction range (dermis) | [0.2,0.3,0.4, 0.7, 1.0, 1.4, 2.1, 3.2, 4.7,7%] |
| Tissue oxygenation | 70% |

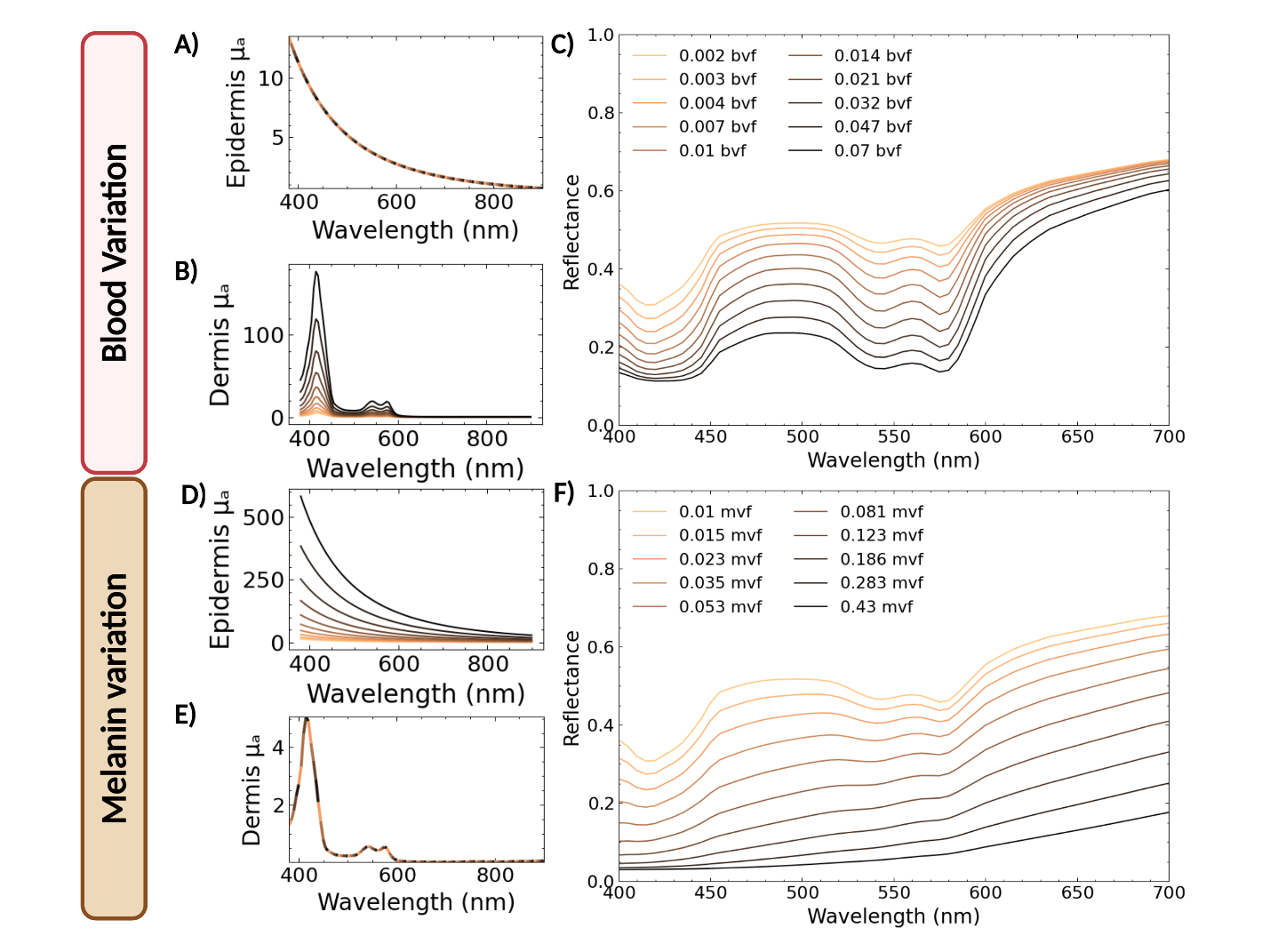

**Figure S1. Model-predicted absorption and reflectance spectra for independent variation of blood and melanin.** Top row (A–C) shows the effects of varying blood volume fraction (bvf) while holding melanosome volume fraction (mvf) constant. (A) Epidermal absorption coefficient µ_a_ (unchanged with bvf), (B) dermal µ_a_ for increasing bvf, and (C) the resulting diffuse reflectance spectra predicted by the adding–doubling (AD) model. Increasing bvf enhances hemoglobin absorption features and reduces reflectance primarily at wavelengths where hemoglobin absorption is strongest. Bottom row (D–F) shows the effects of varying mvf while holding bvf constant. (D) Epidermal µ_a_ for increasing mvf, (E) dermal µ_a_ (unchanged with mvf), and (F) the resulting reflectance spectra. Increasing mvf produces a broadband increase in epidermal absorption and a corresponding decrease in reflectance across the visible spectrum. These reflectance spectra reproduce the expected optical signatures of hemoglobin and melanin in skin.

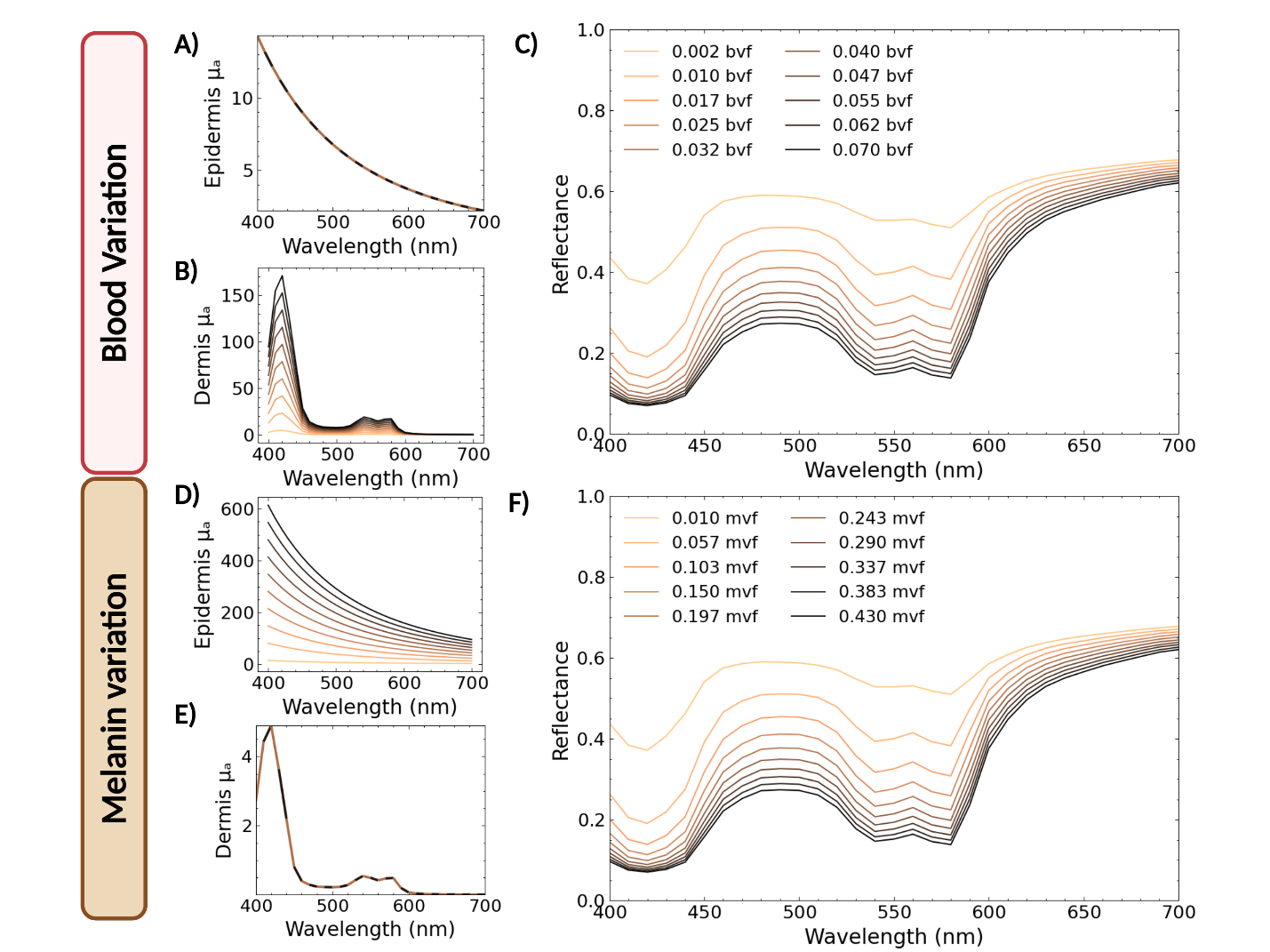

**Figure S2. Monte Carlo model predicted absorption and reflectance spectra for variations of blood and melanin.** (A–C) The effects of varying blood volume fraction (bvf) while holding melanosome volume fraction (mvf) constant. (A) Epidermal absorption coefficient µ_a_ (unchanged with bvf), (B) dermal µ_a_ for increasing bvf, and (C) the resulting diffuse reflectance spectra predicted by the Monte Carlo (MC) model. Increasing bvf enhances hemoglobin absorption features and reduces reflectance primarily at wavelengths where hemoglobin absorption is strongest. Bottom row (D–F) shows the effects of varying mvf while holding bvf constant. (D) Epidermal µ_a_ for increasing mvf, (E) dermal µ_a_ (unchanged with mvf), and (F) the resulting reflectance spectra. Increasing mvf produces a broadband increase in epidermal absorption and a corresponding decrease in reflectance across the visible spectrum. These reflectance spectra reproduce the expected optical signatures of hemoglobin and melanin in skin.

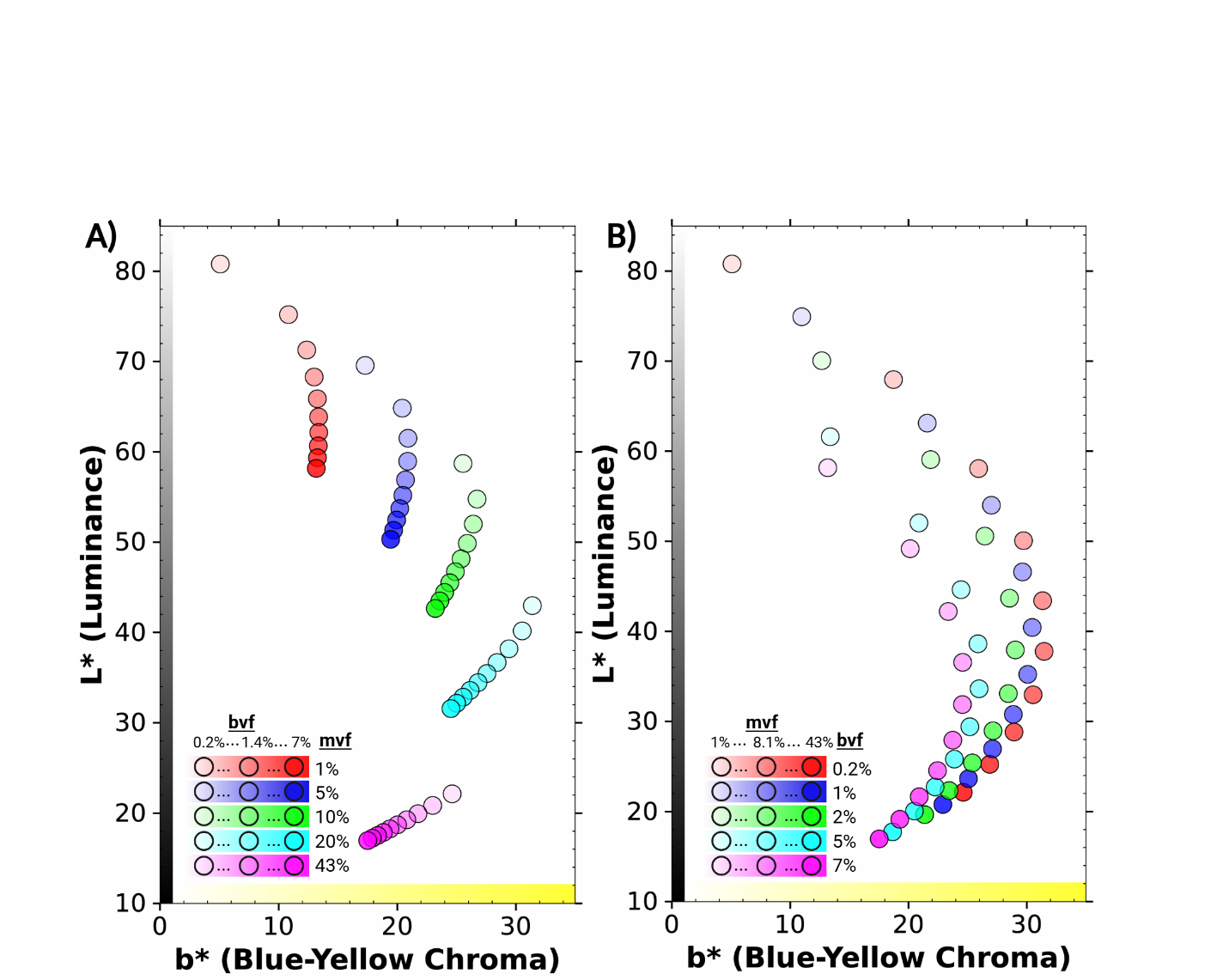

**Figure S3. Iso-melanin and iso-blood trajectories in the L*b* plane generated with the MC model.**  (A) Iso-melanin trajectories in the L*–b* plane generated by holding melanosome volume fraction (mvf) constant while varying blood volume fraction (bvf) using the Monte Carlo (MC) model. Coordinates with same color denote fixed mvf levels, and increasing opacity corresponds to increasing bvf. (B) Iso-blood trajectories generated by holding bvf constant while varying mvf using the MC model. Coordinates with same color denote fixed bvf levels, and increasing opacity corresponds to increasing mvf.

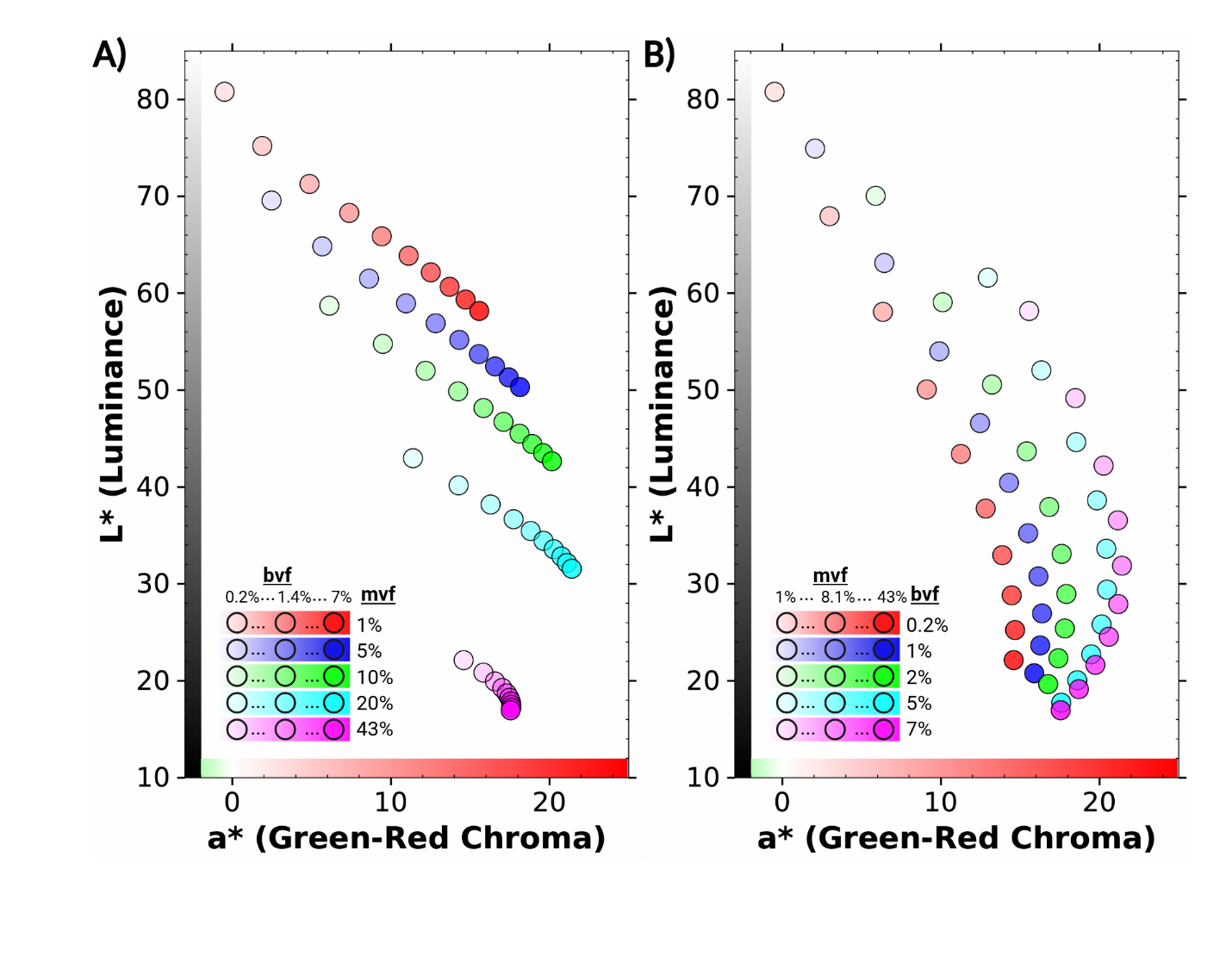

Figure S4. **Iso-melanin and iso-blood trajectories in the L*–a* plane for the MC model.** (A) Iso-melanin trajectories generated by holding melanosome volume fraction (mvf) constant while varying blood volume fraction (bvf) for the MC model. Coordinates with same color denote fixed mvf levels, and increasing opacity corresponds to increasing bvf. Across all mvf levels, increasing bvf produces a decrease in L* and an increase in a*, with the magnitude of this change progressively reduced at higher mvf, resulting in compressed trajectories. (B) Iso-blood trajectories generated by holding bvf constant while varying mvf for the MC model. Coordinates with same color denote fixed bvf levels, and increasing opacity corresponds to increasing mvf. As mvf increases, L* decreases while a* increases and then decreases, producing curved trajectories. As in the L*–b* plane, coordinate spread is largest at low mvf (or low bvf) and progressively decreases at higher mvf, indicating suppression of blood-driven color variation with increasing melanin.

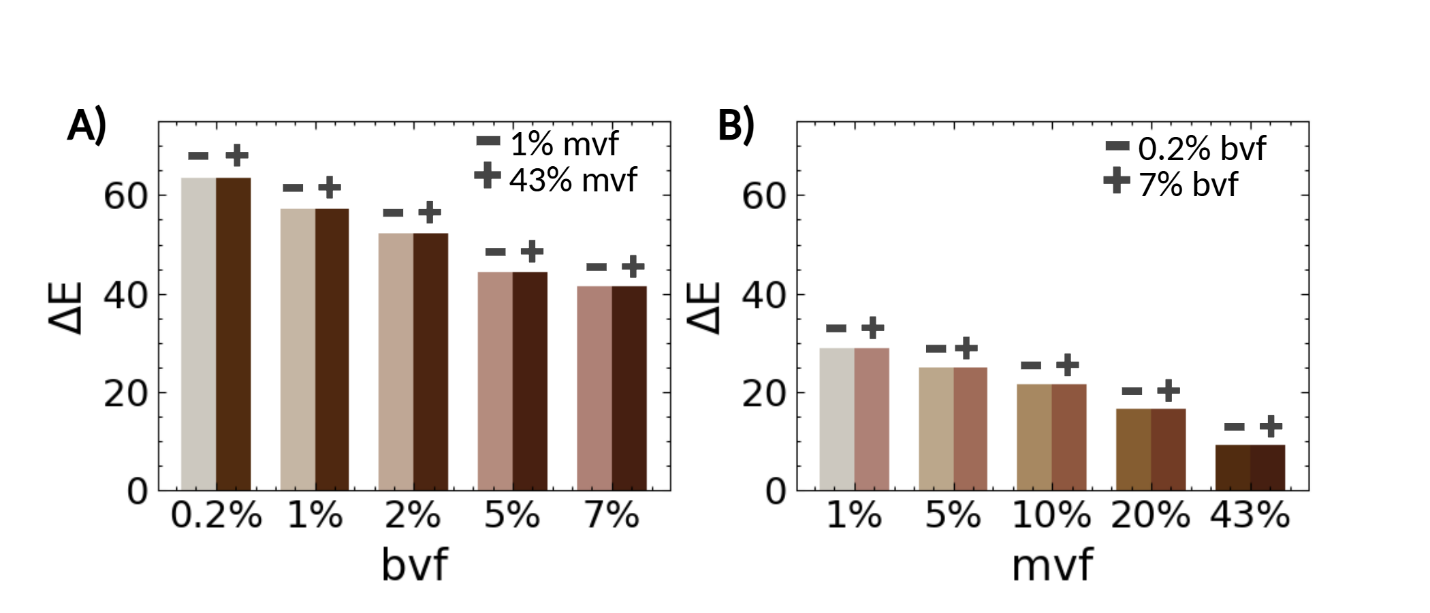

Figure S5. **Perceptible color differences (ΔE) produced by physiological variation in blood and melanin.** (A) ΔE between the minimum and maximum physiological melanosome volume fraction (mvf: 1–43%) at fixed blood volume fraction (bvf) levels predicted by the Monte Carlo (MC) model. (B) ΔE between the minimum and maximum physiological blood volume fraction (bvf: 0.2–7%) at fixed mvf levels predicted by the MC model. Each bar represents the color difference between the L*a*b* coordinates corresponding to the minimum and maximum chromophore values used in the comparison; the two colors indicate the respective L*a*b* values of those endpoints. ΔE associated with variation in blood volume decreases as mvf increases, whereas ΔE associated with variation in melanin remains large across the full range of bvf.

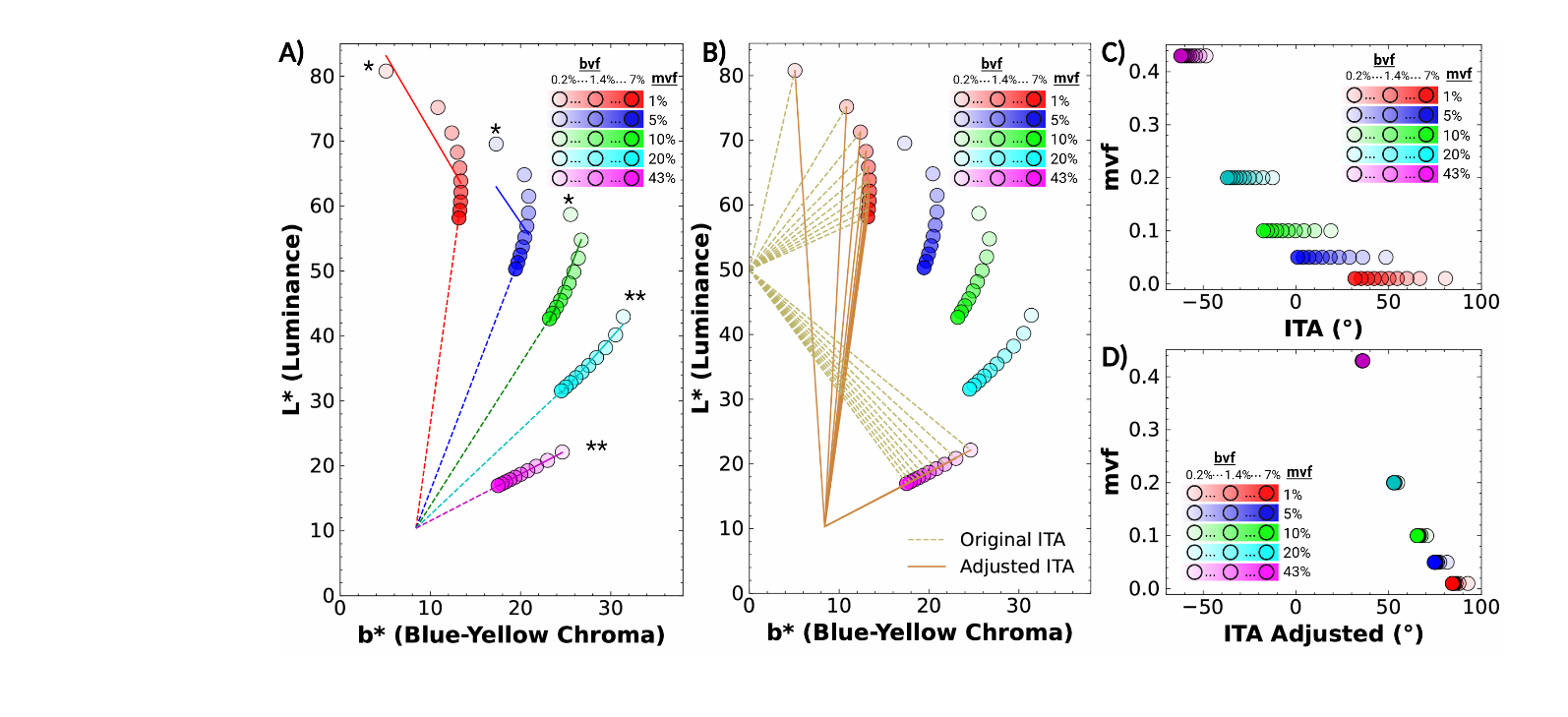

**Figure S6. Adjusted ITA reduces blood-driven variation along MC-model iso-melanin trajectories.** (A) Linear regression of iso melanin trajectories in the L*-b* plane for the MC model. Colored points represent simulated L*–b* coordinates along iso-melanin trajectories generated by varying blood volume fraction (bvf) at fixed melanosome volume fraction (mvf). Solid lines show linear fits across the full trajectories (* R^2^<0.97, ** R^2^≥ 0.97), while dashed lines show fits restricted to the portion of each trajectory where b* decreases with increasing bvf. (B) Geometric interpretation of standard ITA and adjusted ITA. Lines corresponding to constant ITA values are drawn from the standard intercept at (b*, L*) = (0, 50) (solid lines) and from the empirically determined intersection point (b₀, L₀) = (8.4, 10.4) (dashed lines). Examples are shown for the 1% and 43% iso-melanin trajectories. (C) ITA values calculated using the standard formulation (Eq. 1) for all simulated points along iso-melanin trajectories. Substantial variation in ITA occurs along each iso-melanin trajectory, particularly at low mvf levels. (D) ITA values calculated using the adjusted formulation (Eq. 3). The spread of ITA values along iso-melanin trajectories is substantially reduced, particularly at higher mvf levels.

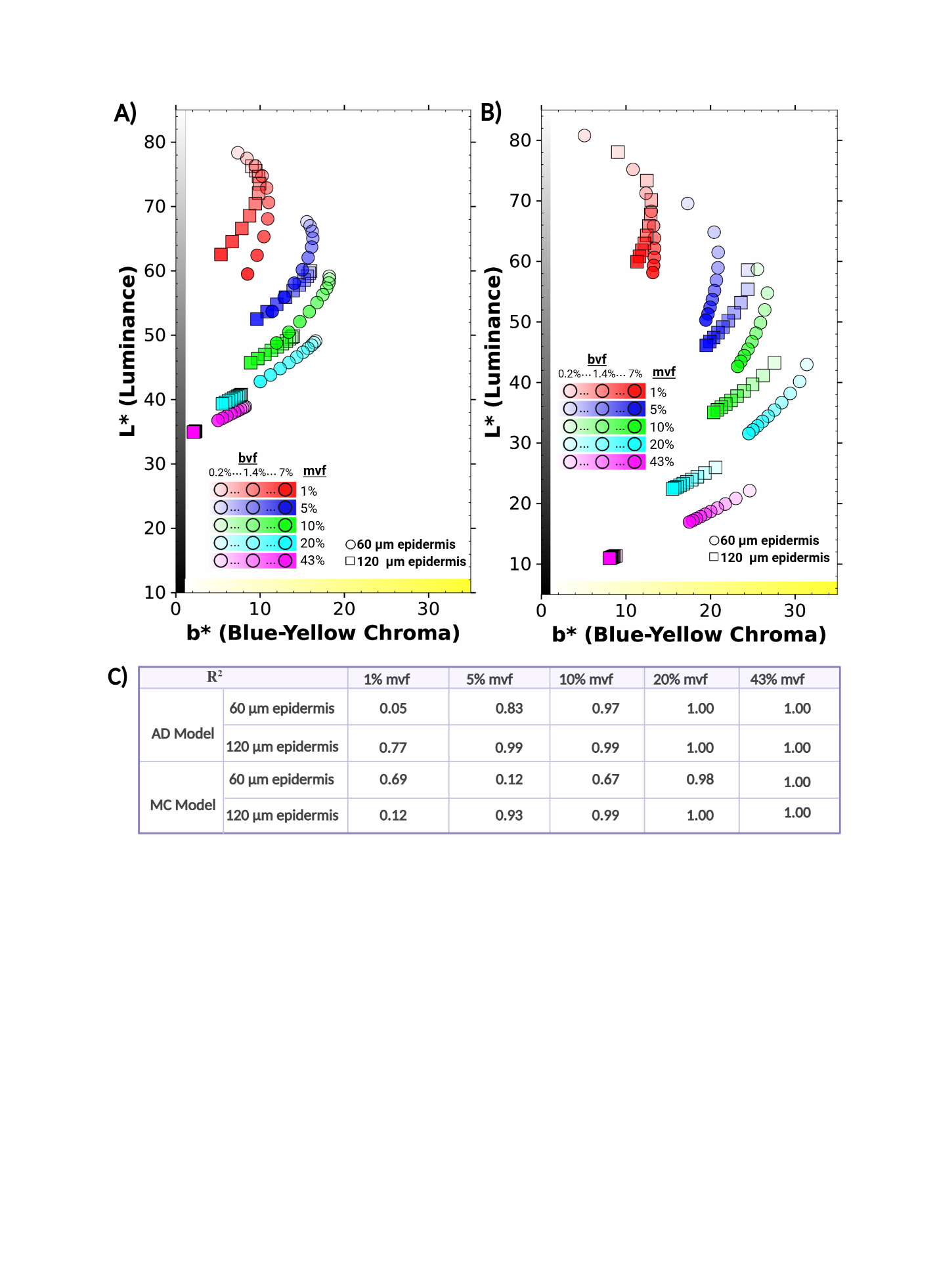

**Figure S7. Epidermal thickness variation preserves high-melanin iso-melanin linearity in the L*–b* plane.** Iso melanin trajectories in the L*-b* plane for varying epidermal thickness with 1 mm dermis using AD model (A) and (B) MC model. Colored points correspond to a given mvf, with increasing opacity indicating increasing bvf. The marker shape denotes the epidermal thickness. (C) The table displays the R^2^ for each trajectory in the L*-b* plane with an epidermal thickness of 120 µm and 60 µm for each model.

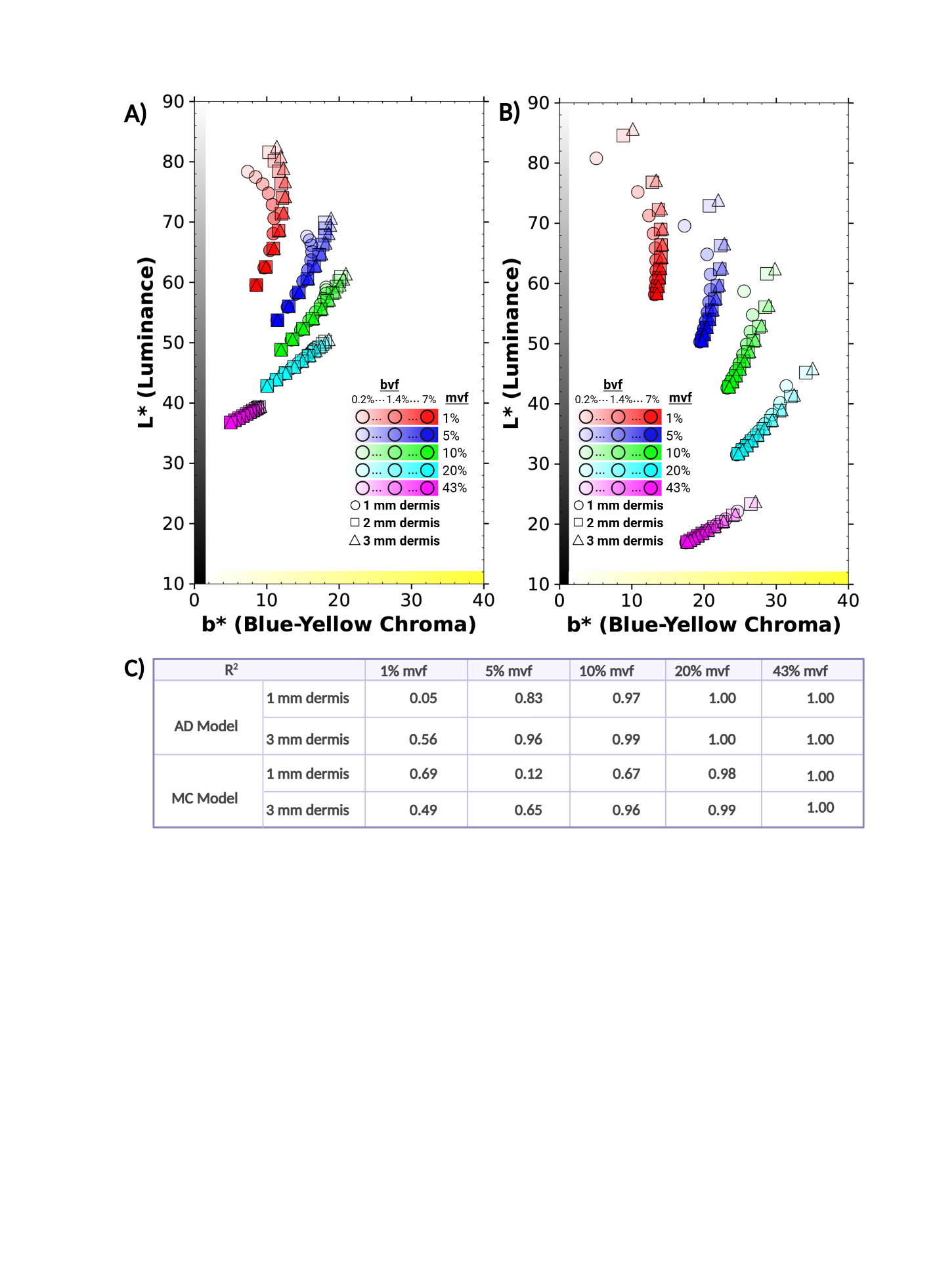

Figure S8. **Dermal thickness variation preserves high-melanin iso-melanin linearity in the L*–b* plane.** Iso melanin trajectories varying dermal thickness with 60 µm epidermis using AD model (A) and (B) using MC model in the L* - b* plane. Colored points correspond to a given mvf, with increasing opacity indicating increasing bvf. The marker shape denotes the dermal thickness. (C) The table displays the R^2^ for each trajectory in the L*-b* plane with a dermal thickness of 1 mm and 3 mm for each model.

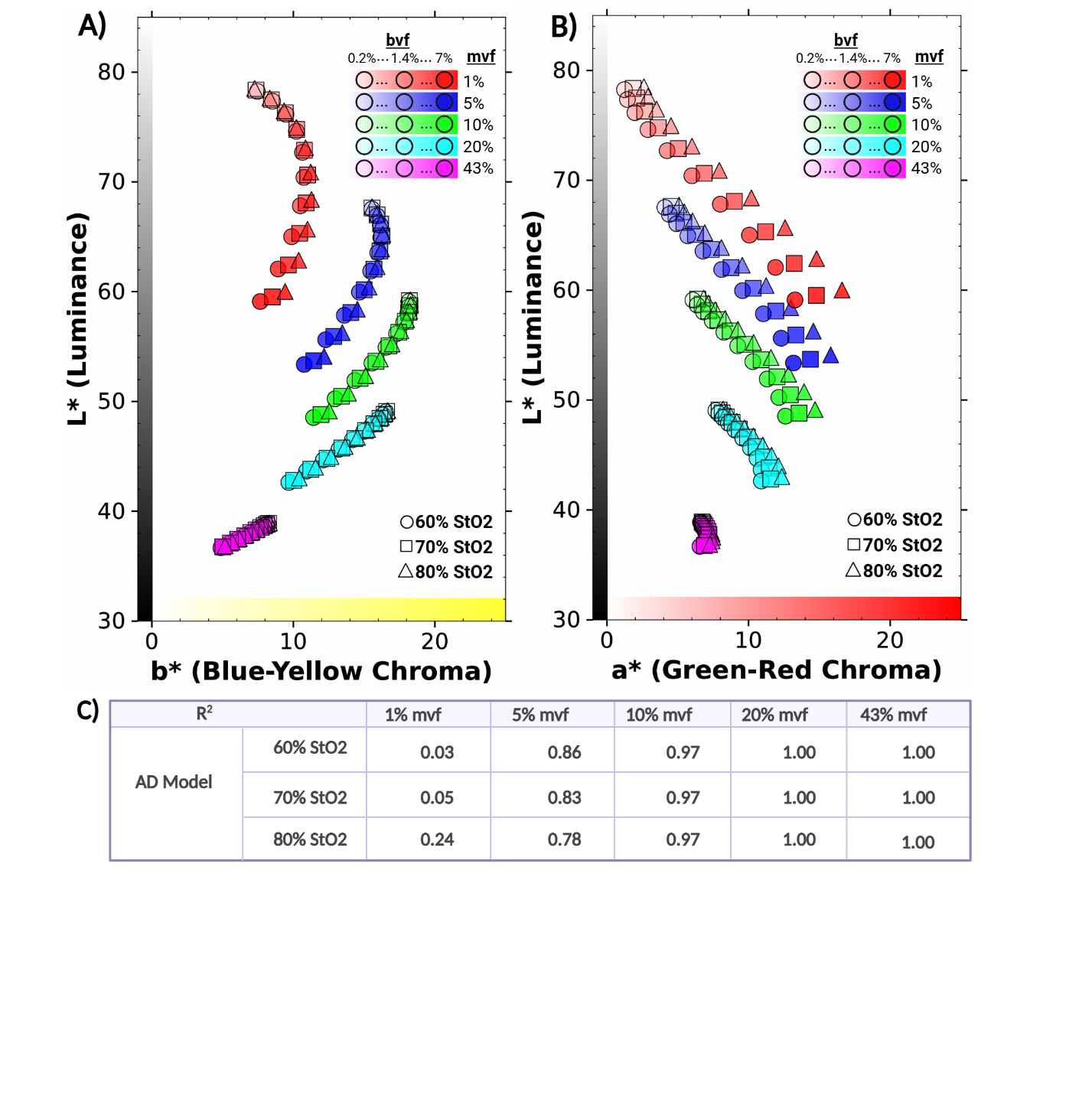

**Figure S9. Tissue oxygenation variation preserves iso-melanin trajectory geometry in the AD model.** Iso melanin trajectories for varying tissue oxygenation using the AD model (A) in the L*-b* plane and (B) in the L*-a* plane. Colored points correspond to a given mvf, with increasing opacity corresponding to increasing bvf. The marker shape denotes the tissue oxygenation levels. (C) The table shows R^2^ values for the iso melanin trajectories in the L*-b* plane. For the lower mvf trajectories (1% and 5%), R^2^ increases as tissue oxygenation increases.

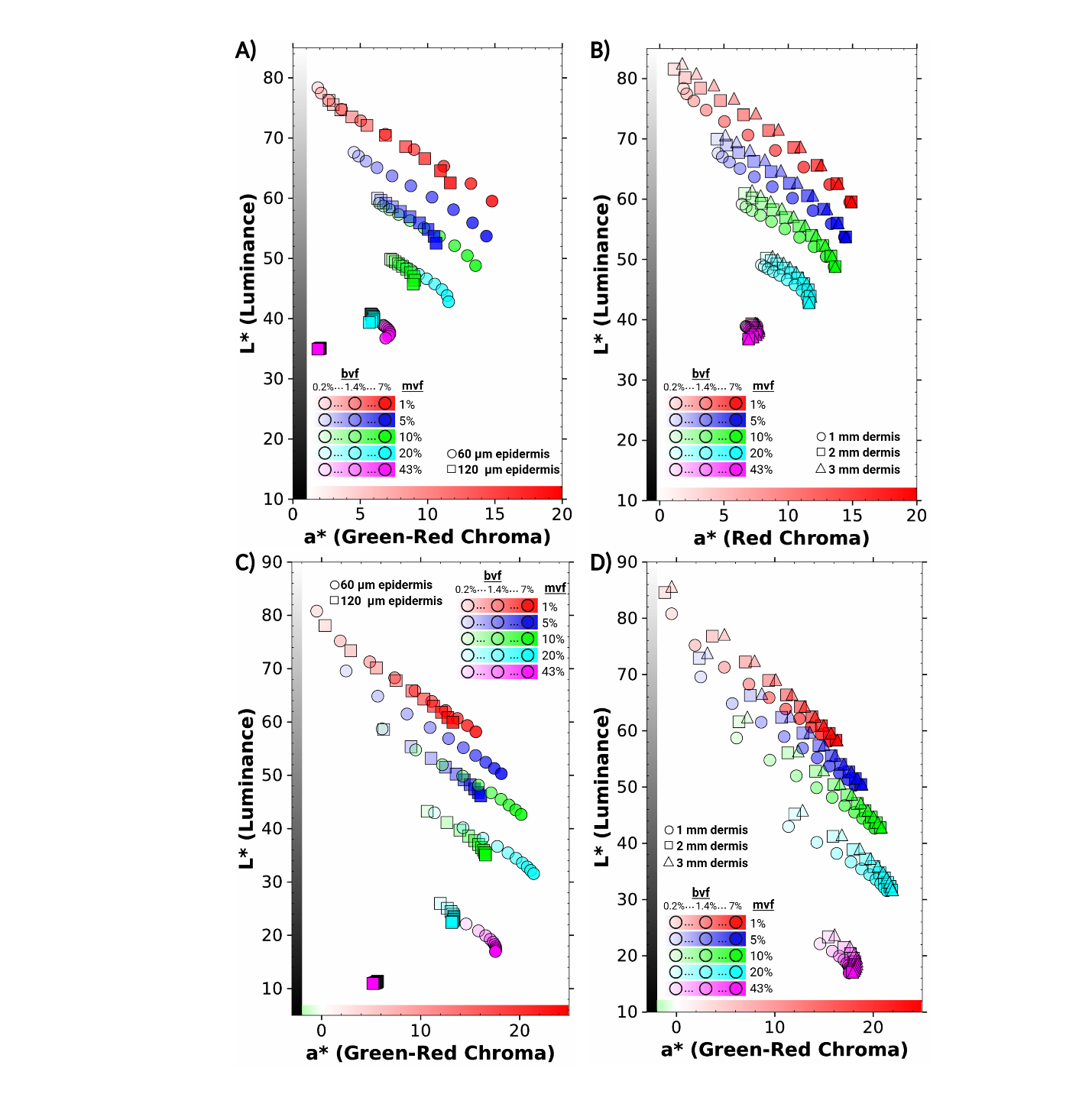

**Figure S10. Iso-melanin trajectory geometry is preserved across epidermal and dermal thickness variation.** Iso melanin trajectories in the L*-a* plane for the AD model when (A) varying epidermal thickness, indicated with marker shape, and (B) when varying dermal thickness, indicated with marker shape. Iso melanin trajectories in the L*-a* plane for the MC model when (C) varying epidermal thickness, indicated with marker shape, and (D) when varying dermal thickness, indicated with marker shape. Colored points correspond to a given mvf, with increasing opacity corresponding to increasing bvf.

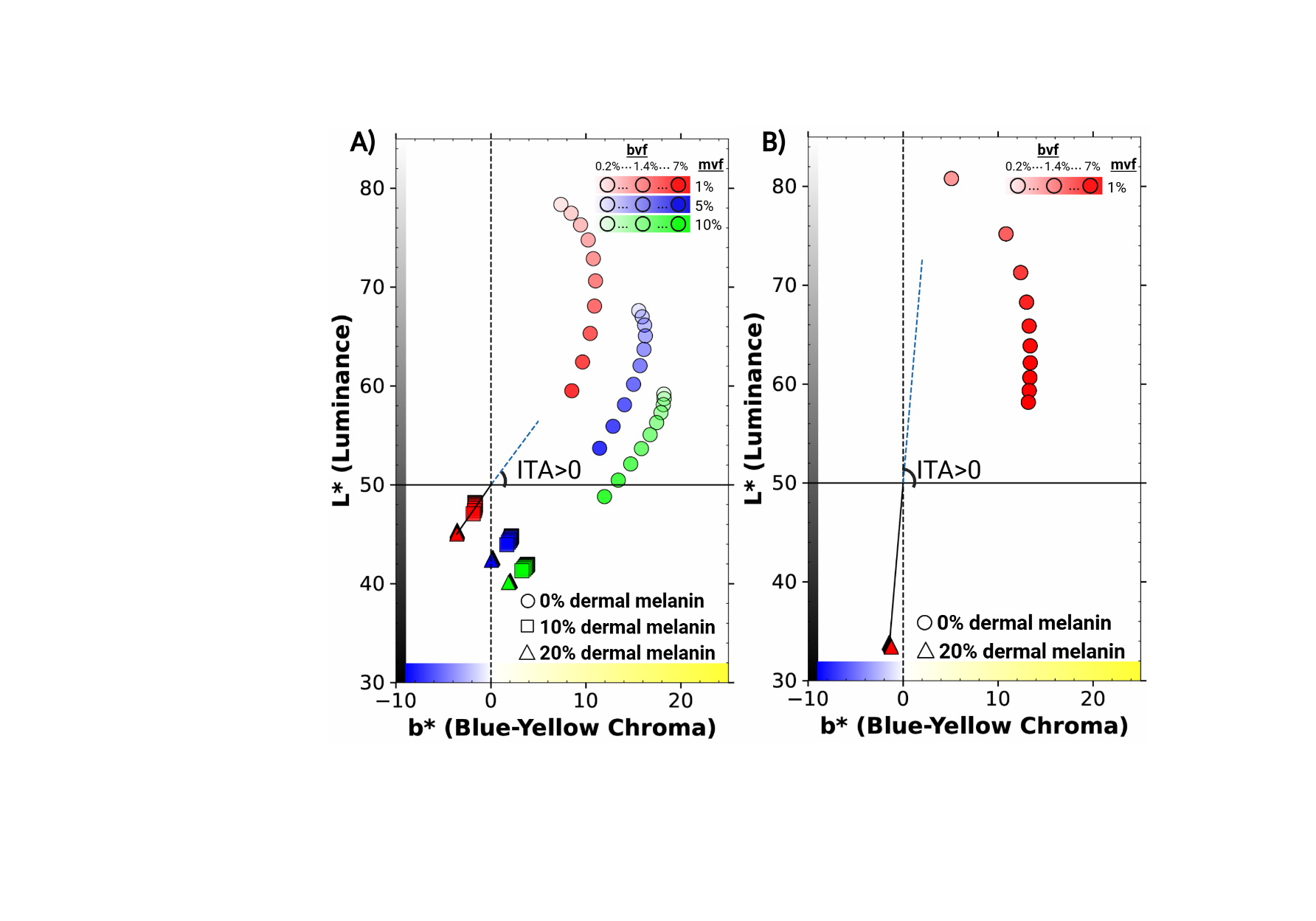

**Figure S11. Dermal melanin shifts skin color into the negative-b* region and produces paradoxically positive ITA values.** Iso melanin trajectories for the AD model in (A) L*-b* with 0%, 10%, and 20% dermal melanin, indicated with marker shape. (B) 1% Iso melanin trajectory for the MC model in the L*-b* plane with 0% and 20% dermal melanin. Colored points correspond to a given mvf, with increasing opacity corresponding to increasing bvf. With increased dermal melanin but lower epidermal melanin content, the b* values become negative, causing positive ITA values.
